# The inner nuclear membrane protein Lem2 coordinates RNA degradation at the nuclear periphery

**DOI:** 10.1101/2021.05.30.446327

**Authors:** Lucía Martín Caballero, Matías Capella, Ramón Ramos Barrales, Nikolay Dobrev, Thomas van Emden, Sabine Fischer-Burkart, Yasuha Kinugasa, Yasuhiro Hirano, Irmgard Sinning, Tamás Fischer, Yasushi Hiraoka, Sigurd Braun

**Affiliations:** BioMedical Center (BMC), Division of Physiological Chemistry, Faculty of Medicine, LMU Munich, Planegg-Martinsried, Germany; International Max Planck Research School for Molecular and Cellular Life Sciences, Planegg-Martinsried, Germany; Centro Andaluz de Biología del Desarrollo (CABD), Universidad Pablo de Olavide-Consejo Superior de Investigaciones Científicas-Junta de Andalucía, Ctra. Utrera km.1, Seville, Spain; Heidelberg University Biochemistry Center (BZH), Heidelberg, Germany; European Molecular Biology Laboratory, Hamburg Unit, Hamburg, Germany; Graduate School of Frontier Biosciences, Osaka University, Suita, Japan; The John Curtin School of Medical Research, The Australian National University, Canberra, Australia; Research Institute for Radiation Biology and Medicine, Hiroshima University, 1-2-3 Kasumi, Minamiku, Hiroshima 734-8553, Japan

**Keywords:** LEM-containing proteins, nuclear exosome, RNA surveillance, non-coding RNA, meiosis

## Abstract

Transcriptionally silent chromatin often localizes to the nuclear periphery. However, whether the nuclear envelope (NE) is a site for post-transcriptional gene repression is unknown. Here we demonstrate that S. pombe Lem2, an NE protein, regulates nuclear exosome-mediated RNA degradation. Lem2 deletion causes accumulation of non-coding RNAs and meiotic transcripts. Indeed, an engineered exosome substrate RNA shows Lem2-dependent localization to the nuclear periphery. Lem2 does not directly bind RNA, but instead physically interacts with the exosome-targeting MTREC complex and promotes RNA recruitment. The Lem2-assisted pathway acts independently of nuclear bodies where exosome factors assemble, revealing that multiple spatially distinct degradation pathways exist. The Lem2 pathway is environmentally responsive: nutrient availability modulates Lem2 regulation of meiotic transcripts. Our data indicate that Lem2 recruits exosome co-factors to the nuclear periphery to coordinate RNA surveillance and regulates transcripts during the mitosis-to-meiosis switch.

---

Eukaryotic genomes are pervasively transcribed, giving rise to sense and antisense RNAs from intra- and intergenic regions and repetitive elements. This widespread transcription is ubiquitously found across lower eukaryotes and metazoans (Jensen et al., 2013). Accumulation of cryptic unstable transcripts (CUTs) may cause genome instability through RNA-DNA hybridization, posing a general threat to cells (Wahba et al., 2013). Other coding and non-coding RNAs are continuously transcribed but only function during specific developmental stages. A prominent example is transcription of meiotic mRNAs in fission yeast, whose expression is temporally restricted by fast turnover in vegetative cells (Harigaya et al., 2006; Hiriart et al., 2012; Tashiro et al., 2013; Zofall et al., 2012). Constant surveillance is thus required to prevent the aberrant or untimely accumulation of these different types of transcripts. Such regulation is mediated by a variety of nuclear and cytosolic pathways that are controlled by endogenous and environmental factors. Yet how these different RNA degradation processes are coordinated is poorly understood. In particular, while transcriptional silencing has been intensely studied in the context of perinuclear anchoring, which facilitates local concentration of silencing factors, the role of the NE in post-transcriptional repression remains largely unknown.

Nuclear RNA degradation in eukaryotes is mediated by the nuclear exosome, a barrel-shaped multiprotein complex with a central channel and two associated 3’-5’ exoribonucleases, Rrp6 and Dis3/Rrp44 (Schmid and Jensen, 2019). RNA surveillance is assisted by exosome-targeting complexes conserved across eukaryotes, which contribute to substrate specificity and RNA processing (Schmid and Jensen, 2019). They contain polyadenylation and RNA helicase activities that facilitate unwinding of RNA secondary structures to feed substrates into the exosome channel. In the fission yeast *Schizosaccharomyces pombe*, these complexes include TRAMP and MTREC. TRAMP (<u>Tr</u>f4/5-<u>A</u>ir1/2-<u>M</u>tr4 <u>p</u>olyadenylation) consists of a non-canonical poly(A) polymerase (Cid14; Trf4/5 in *S. cerevisiae*), a zinc-knuckle RNA-binding protein (Air1), and an RNA helicase (Mtr4). Fission yeast TRAMP contributes to small nucleolar RNA (snoRNA) processing as well as the degradation of antisense transcripts and RNAs derived from pericentromeric repeats (Bühler et al., 2008; Zhang et al., 2011; Zhou et al., 2015). The MTREC (<u>M</u>tl1-<u>Re</u>d1 <u>c</u>ore) complex comprises the Mtr4-like helicase Mtl1 and the zinc-finger domain protein Red1 (<u>R</u>NA <u>e</u>limination <u>d</u>efective 1). This complex, also known as NURS (<u>nu</u>clear <u>R</u>NA <u>s</u>ilencing) complex, mediates the turnover of CUTs, meiotic and non-spliced transcripts (Egan et al., 2014; Lee et al., 2013; Zhou et al., 2015) and is orthologous to the human PAXT (<u>p</u>oly(<u>A</u>) tail e<u>x</u>osome <u>t</u>argeting) complex (Ogami et al., 2018). MTREC assembles into an 11-subunit “super complex”, in which Red1 acts as a central scaffold for different submodules and makes contact with Rrp6 via Mtl1 (Egan et al., 2014; Lee et al., 2013; Sugiyama and Sugioka-Sugiyama, 2011; Zhou et al., 2015). The submodules have different activities and include the canonical Poly(A) polymerase Pla1 and the complexes Red5-Pab2-Rmn1, Ars2-Cbc1-Cbc2 complex, and Iss10-Mmi1 (Dobrev et al. *in press*; Lemay et al., 2010; Lemieux et al., 2011; St-André et al., 2010; Sugiyama et al., 2013; Thillainadesan et al., 2020; Yamanaka et al., 2010; Zhou et al., 2015).

Recognition and degradation of exosome targets in *S. pombe* is best understood in the context of meiotic transcript turnover. The YTH (<u>YT</u>521-B <u>h</u>omology) protein Mmi1 recognizes hexanucleotide (UNAAAC) motifs known as DSR (<u>d</u>eterminant of <u>s</u>elective <u>r</u>emoval) sequences, which are present in meiotic transcripts (Harigaya et al., 2006; Hiriart et al., 2012; Yamashita et al., 2012). Similar motifs are linked to intron retention detection in several pre-mRNAs (Kilchert et al., 2015). Substrate binding requires Mmi1 dimerization and interaction with its partner Erh1 (<u>e</u>nhancer of <u>r</u>udimentary <u>h</u>omolog 1), resulting in the formation of the tetrameric <u>E</u>rh1-<u>M</u>mi1 <u>c</u>omplex (EMC). EMC sequesters RNA substrates, preventing nuclear export and translation (Andrić et al., 2021; Hazra et al., 2020; Shichino et al., 2018; Sugiyama et al., 2016; Xie et al., 2019). Mmi1 further associates with the Ser-/Pro-rich protein Iss10 (also known as Pir1), which bridges Mmi1-MTREC interaction (Lee et al., 2013; Sugiyama and Sugioka-Sugiyama, 2011; Yamashita et al., 2013; Zhou et al., 2015). Several DSR-containing genes are also marked by histone H3 lysine 9 methylation (H3K9me) in an Mmi1-, Red1- and Rrp6-dependent manner, potentially adding another layer of control to prevent ectopic expression of these meiotic transcripts (Egan et al., 2014; Zofall et al., 2012). At other exosome-targeted genes, Rrp6 inactivation or stress exposure induces an alternative degradation pathway employing RNA interference (RNAi), which also results in deposition of H3K9me at several loci known as HOODs (<u>h</u>eter<u>o</u>chr<u>o</u>matin <u>d</u>omains) (Yamanaka et al., 2013).

The exosome, and its co-factors, assemble into one or several nuclear foci in mitotically growing cells. The presence of EMC-containing nuclear foci depends on Mmi1 dimerization and its association with Erh1 (Shichino et al., 2020; Sugiyama et al., 2016), whereas Iss10 is critical for Red1 foci formation and the co-localization of MTREC with EMC (Shichino et al., 2020; Sugiyama and Sugioka-Sugiyama, 2011; Yamashita et al., 2013). In vegetative cells, EMC also associates with the CCR4-NOT complex and a ncRNA known as *mamRNA* to counterbalance its own inhibitor, Mei2, through ubiquitin-dependent degradation (Andrić et al., 2021; Cotobal et al., 2015; Simonetti et al., 2017; Stowell et al., 2016; Sugiyama et al., 2016; Ukleja et al., 2016). Upon the onset of meiosis, Iss10 becomes unstable and Red1 foci disappear (Sugiyama and Sugioka-Sugiyama, 2011; Wei et al., 2021). This causes the dissociation of Mmi1 from MTREC and its collapse into a single focus known as the Mei2 dot (Harigaya et al., 2006; Shichino et al., 2014). The Mei2 dot forms a nuclear structure comprising the RNA-binding protein Mei2 and the ncRNA *sme2* (also known as *meiRNA*). Together, Mei2 and *meiRNA* sequester Mmi1 at the genomic locus of *sme2*^+^, resulting in the inactivation of Mmi1-dependent RNA elimination (Ding et al., 2012; Shichino et al., 2014; Shimada et al., 2003). Iss10 degradation and the disassembly of nuclear foci coincide with the accumulation of meiotic transcripts. It has therefore been proposed that Iss10-dependent nuclear foci present specific sites of RNA degradation. However, cells lacking Iss10 or expressing a mutant deficient in Iss10-Red1 interaction (*red1*-Δ*196-347*) display only minor defects in RNA turnover (Egan et al., 2014; Shichino et al., 2020; Yamashita et al., 2013); conversely, a Red1 point mutant deficient in RNA degradation (*red1-H637I*) does not affect its ability to form nuclear foci (Sugiyama and Sugioka-Sugiyama, 2011). Hence, the physiological function of these nuclear foci remains poorly understood, and whether other nuclear structures contribute to RNA turnover has not been addressed.

Transcriptionally silent chromatin and repressive histone modifying enzymes are often sequestered at the nuclear periphery, which is thought to promote nucleation and spreading of heterochromatin (Harr et al., 2016). In *S. pombe*, perinuclear heterochromatin is marked by H3K9me, which is deposited by the histone H3K9 methyltransferase Clr4 (Nakayama et al., 2001) and recognized by HP1 (<u>h</u>eterochromatin <u>p</u>rotein <u>1</u>) family chromodomain proteins (Bannister et al., 2001; Sadaie et al., 2008) This heterochromatic platform serves as a recruitment site for the <u>S</u>nf2-like/<u>H</u>DAC-containing <u>re</u>pressor <u>c</u>omplex SHREC, restricting access to RNA polymerase II (Pol II) (Sugiyama et al., 2007). Several NE proteins contribute to the proper silencing and perinuclear localization of heterochromatin in *S. pombe* (Barrales et al., 2016; Holla et al., 2020; Iglesias et al., 2020). Lem2 is a conserved integral protein of the inner nuclear membrane (INM) with two nucleoplasmic domains, the LEM (<u>L</u>AP2, <u>e</u>merin, <u>M</u>AN1) and MSC (<u>M</u>AN1-<u>S</u>rc1p <u>C</u>-terminal) domain. While the LEM domain contributes to centromere tethering, the MSC domain is important for telomere anchoring and transcriptional silencing of heterochromatin (Banday et al., 2016; Barrales et al., 2016; Hirano et al., 2018; Tange et al., 2016). Lem2 acts in the same silencing pathway as SHREC and promotes its recruitment to heterochromatin, providing a functional link between heterochromatin silencing and nuclear organization (Barrales et al., 2016). Whether Lem2 contributes to other modes of gene regulation remains unknown.

Here, we uncover a global role for Lem2 in repressing non-coding RNAs and meiotic genes, which is distinct from its function in heterochromatin silencing. We show that Lem2 cooperates with the nuclear exosome and physically interacts with the MTREC subunit Red1. Importantly, we demonstrate that Lem2 is critical for the perinuclear localization of exosome substrates and their recognition by Mmi1 and Red1. Lem2-assisted RNA targeting is independent of, and partially redundant with, the degradative functions of Iss10 and Erh1. In addition, Lem2 contributes to CUT degradation. This implies that multiple RNA degradation pathways exist and localize to different subnuclear structures. Finally, we show that Lem2 activity is regulated by nutrient availability, and is inactivated upon starvation in parallel to Iss10 degradation. We propose that Lem2 assists in RNA surveillance by coordinating the degradation of exosome targets at the nuclear periphery. Lem2 is therefore mechanistically essential for fine-tuning the gene expression program during meiotic onset through nutrient-dependent regulation.

## Results

### Lem2 mediates repression of non-coding RNAs and meiotic genes

The inner nuclear membrane protein Lem2 plays a crucial role in tethering and silencing of constitutive heterochromatin (Barrales et al., 2016; Tange et al., 2016). To examine whether Lem2 regulates gene expression through additional mechanisms, we performed transcriptome analysis using whole-genome RNA sequencing (RNA-seq) in wild-type (WT) and *lem2*Δ mutant cells. We found that a significant portion of the genome was upregulated in *lem2*Δ cells, whereas only a few transcripts were decreased (838 vs. 35, from a total of 6642 transcripts; Fig. 1A,B). The *S. pombe* genome contains roughly 70% protein coding and 21% non-coding genes (Lock et al., 2019). Remarkably, 61% of upregulated transcripts in *lem2*Δ cells were non-coding RNAs (ncRNAs), indicating significant over-representation of this transcript type (Fig. 1A,C). Upregulated ncRNAs include the meiotic transcript *sme2*^+^, which plays a key role in meiotic onset in *S. pombe* (Watanabe and Yamamoto, 1994) and the small nucleolar RNA *sno20*^+^ (Fig. 1D). In addition, we found several long-terminal repeats (LTRs) among the most upregulated transcripts (Fig. 1A,C). Transcript upregulation was restricted to individual loci, as the expression of neighboring genes remained unaffected (Fig. 1D).

**Fig. 1:**
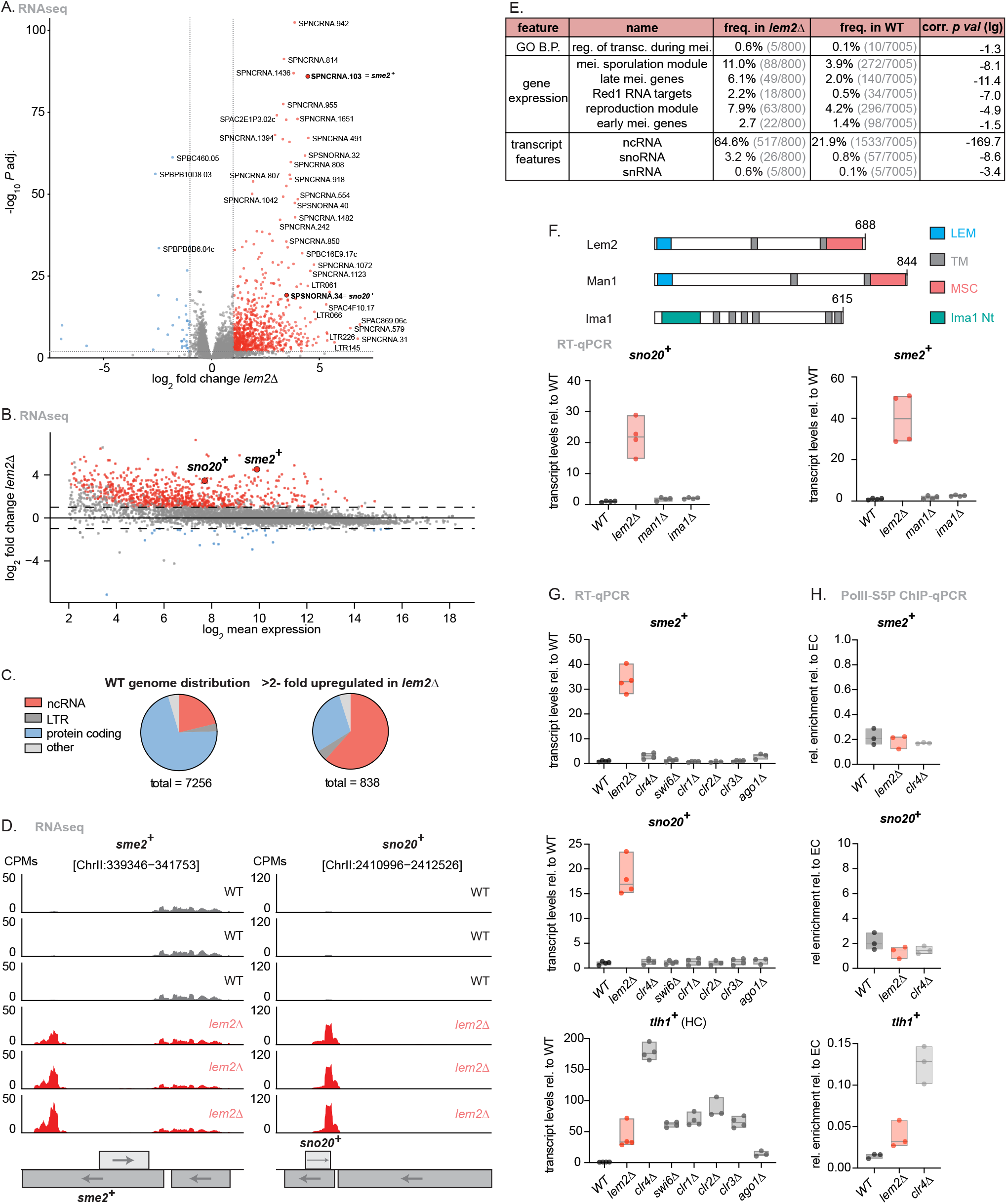
Lem2 represses non-coding RNAs and meiotic genes. **A.** RNAseq data of *lem2*Δ vs WT displayed as a volcano plot depicting statistical significance (Y axis) versus fold change (X axis). Significant (−log_10_ *p* adj. value > 2) genes are highlighted, both upregulated (log_2_ fold change > 1 in red) and downregulated (log_2_ fold change < −1 in blue). Some of the most upregulated transcripts are highlighted. **B.** MA-Plot of *lem2*Δ cells relative to WT. X axis shows the log_2_ mean expression and the Y axis shows the log_2_ fold change in *lem2*Δ over WT. **C.** Pie charts showing transcript feature distributions (ncRNA, LTR, protein coding, other (pseudogene, rRNA, snoRNA, snRNA, tRNA). Left: genome-wide distribution of transcript features in a WT genome. Right: transcript feature distribution of the significantly upregulated (log_2_ fold change > 1, −log_10_ *p* adj. value > 2) transcripts in *lem2*Δ cells. **D.** Coverage plots showing upregulated transcripts in *lem2*Δ cells. The Y axis shows normalized read counts (counts per million (CPMs)). Three independent biological replicates are shown per condition. Genomic coordinates are shown in base pairs (bp) above the graphs. **E.** Table with selected results from gene list enrichment analysis of *lem2*Δ mutants. The Bähler Lab *AnGeLi* tool with FDR=0.05 was used for this analysis (Bitton et al., 2015). GO B.P. = Gene Ontology Biological Process; reg. of transc. during mei. = regulation of transcription during meiosis; mei. =meiotic. **F.** Top: Domain structure of INM proteins Lem2, Man1 and Ima1. Protein length is given in amino acids. Bottom: Transcript levels of *sme2*^+^ and *sno20*^+^ quantified by RT-qPCR on the indicated strains. **G.** *sme2^+^, sno20*^+^ and *tlh1*^+^ transcript levels quantified by RT-qPCR in the indicated strains. **H.** ChIP-qPCR analysis of Pol II-S5P enrichment of *sme2^+^, sno20*^+^ and *tlh1*^+^ in the indicated strains. The data was divided by input and normalized to euchromatin levels (*act1*^+^, *tef3*^+^, *ade2*^+^). n = 3 independent biological replicates. For F and G, data normalized to *act1*^+^ expression is shown relative to WT. n = 3-4 independent biological replicates. For F, G and H, the individual replicates are shown in a floating bar plot and the line depicts the median.

We further examined the upregulated transcripts using the Analysis of Gene Lists (AnGeLi) online tool for comprehensive interrogation of gene lists in S. pombe (Bitton et al., 2015). This analysis revealed “ncRNA” as the most significant group. In addition, we found several other genes linked to early and late meiosis (Fig. 1E; a complete list of features can be found in the Suppl. Table 1). We examined transcript levels in mutants deficient for two other integral envelope proteins known to interact with chromatin, Man1 and Ima1 (Steglich et al., 2012). Genome-wide analysis of man1Δ cells revealed no major transcriptional changes (Suppl. Fig. 1A). Using reverse transcription followed by quantitative PCR (RT-qPCR), we confirmed the upregulation of selected meiotic and non-coding RNA transcripts in the lem2Δ strain; in contrast, these transcripts were largely unaltered in man1Δ and ima1Δ cells (Fig. 1F, Suppl. Fig. 1D). This indicates that Lem2 is specifically associated with regulation of these transcripts.

We previously showed that Lem2 participates in transcriptional silencing by recruiting the repressor complex SHREC to pericentromeric and subtelomeric heterochromatin (Barrales et al., 2016). We therefore examined whether heterochromatin factors also regulate meiotic transcripts and ncRNA expression. By performing RNA-seq in cells lacking the H3K9 methyltransferase Clr4, we found increased levels of pericentromeric ncRNAs and subtelomeric mRNAs as previously shown (Cam et al., 2005; Gallagher et al., 2018) (Suppl. Fig. 1B). While many of the transcripts upregulated in *clr4*Δ cells are also increased in cells lacking Lem2, in agreement with its role in heterochromatin silencing, the overall number of upregulated transcripts was significantly lower in the *clr4*Δ mutant (103 vs. 838, Suppl. Fig. 1B,C). We examined selected targets by RT-qPCR in *clr4*Δ and mutants lacking other factors involved in heterochromatin regulation. The processes we examined include heterochromatin spreading (*swi6*Δ), RNAi (*ago1*Δ), and restriction of RNA polymerase II (Pol II) access by the SHREC complex (*clr1*Δ, *clr2*Δ, *clr3*Δ). In stark contrast to heterochromatic genes, *sme2*^+^ and *sno20*^+^ were unaltered in these heterochromatin mutants (Fig. 1G, Suppl. Fig. 1E). *ssm4*^+^ showed modest upregulation in *clr4*Δ and *ago1*Δ mutants (Suppl. Fig. 1E), in agreement with its location within a heterochromatin island (Zofall et al., 2012). These data suggest that Lem2 regulates the expression of these exosome targets largely independently of heterochromatin formation.

While heterochromatic loci are controlled at the transcriptional and post-transcriptional levels, meiotic genes, including *sme2*^+^, are mainly regulated through RNA degradation by the nuclear exosome (Harigaya et al., 2006; Hiriart et al., 2012; Yamamoto, 2010; Yamashita et al., 2012). To determine whether Lem2 acts at the transcriptional or post-transcriptional level, we performed chromatin immunoprecipitation followed by qPCR (ChIP-qPCR) with the elongating form of RNA polymerase II, which is phosphorylated at serine 5 in its C-terminal domain (CTD) (Pol II-S5P). As expected, *clr4*Δ cells showed strong enrichment of Pol II-S5P at *tlh1*^+^, a subtelomeric, heterochromatic locus (Fig. 1H), which correlated with increased transcript levels (Fig. 1G). We also observed moderate Pol II-S5P enrichment at *tlh1*^+^ in cells lacking Lem2, consistent with its role in heterochromatin (Barrales et al., 2016). However, Pol II-S5P abundance was unaltered at the *sme2*^+^ and *sno20*^+^ genes in *lem2*Δ cells (Fig. 1H), despite the increased transcript levels (Fig. 1G). These results further corroborate the notion that Lem2 regulates meiotic and non-coding transcripts through a mechanism that is likely transcription-independent and distinct from its previously described role in heterochromatin regulation.

### Lem2 cooperates with the nuclear exosome in nuclear RNA surveillance

As meiotic transcripts and ncRNAs are major targets of the nuclear exosome (Zhou et al., 2015) (Fig. 2A), we examined whether the post-transcriptional role of Lem2 is linked to nuclear RNA degradation. We performed RNA-seq with several mutants lacking components of the nuclear exosome pathway: the 3’-5’ exoribonuclease Rrp6 (nuclear exosome); the zinc-finger protein Red1 (MTREC); Erh1 (EMC complex); the 3’-5’ exoribonuclease Ccr4 (CCR4-NOT complex); the zinc-finger protein Air1 (TRAMP) (see Fig. 2A). Principal component analysis (PCA) revealed mutant-specific groups displaying high reproducibility across most independent biological replicates (Suppl. Fig. 2A). We conducted differential expression analysis followed by unsupervised K-means clustering and observed a striking similarity for the transcriptome profiles of *lem2*Δ, *rrp6*Δ, and *red1*Δ mutants, whereas *lem2*Δ showed only limited overlap with *air1*Δ, *erh1*Δ and *ccr4*Δ (Fig. 2B). Pairwise transcriptome comparison revealed a strong positive correlation for *lem2*Δ with *rrp6*Δ and *red1*Δ (R = 0.79 and 0.65, respectively; Fig. 2C, Suppl. Fig. 2B) and a weaker correlation with *air*Δ (R = 0.53, Suppl. Fig. 2C); conversely, no correlation was seen with *erh1*Δ or *ccr4*Δ at the genome-wide level (Suppl. Fig. 2C). We also found a strong accumulation of anti-sense transcripts in the *lem2*Δ strain similar to *rrp6*Δ (Suppl. Fig. 2D). Using *AnGeLi* (Bitton et al., 2015), we next analyzed the top clusters containing the upregulated transcripts in *lem2*Δ, *rrp6*Δ and *red1*Δ strains (clusters 1-5). Cluster 1 was enriched for features related to early meiosis (38% frequency) and Red1-mediated degradation (24%), and many of these transcripts were also increased in cells lacking Erh1, consistent with its role in binding to these exosome substrates as part of EMC (Fig. 2B, Suppl. Fig. 2E). In contrast, transcripts present in clusters 2, 3, 4 and 5 were predominantly enriched for ncRNAs (59%, 63%, 45% and 73%, respectively), RNA splicing, or genes related to stress regulation. For cluster 2, we also observed specific overlap with the *air1*Δ mutant (Fig. 2B). In addition, several middle and late meiotic genes and transcripts related to sporulation were present in cluster 3 and 4 (Fig. 2B, Suppl. Fig. 2E; a complete list of features is provided in Suppl. Table 2). Together, these results suggest that Lem2 cooperates with distinct exosome factors to control transcripts through multiple degradation pathways.

**Fig. 2:**
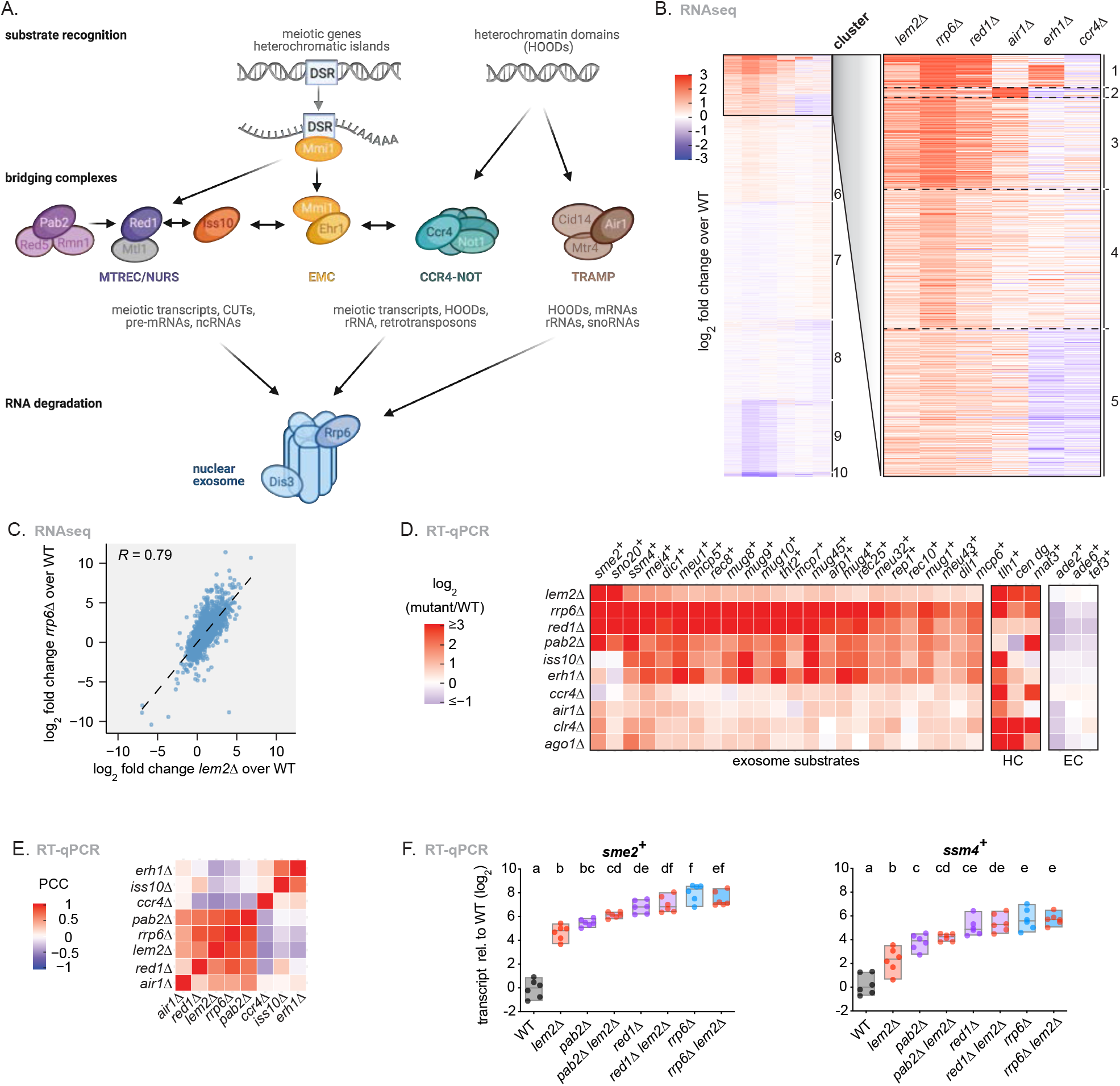
Lem2 cooperates with the nuclear exosome. **A.** Simplified scheme highlighting some of the main players in the nuclear exosome pathway. The core nuclear exosome contains subunits with exoribonucleic catalytic activity. Different targeting/bridging complexes (MTREC/NURS, EMC, CCR4-NOT and TRAMP) provide substrate specificity. YTH protein Mmi1 can directly bind DSR motifs in meiotic genes. Heterochromatic domains (HOODs) are partially controlled via CCR4-NOT and TRAMP. **B.** K-means clustering of RNAseq data. Genes were clustered using K-means clustering based on their differential expression in the indicated mutants. Ten clusters were generated and the clusters with most upregulated transcripts in *lem2*Δ (clusters 1-5) were analyzed with *AnGeLi* with FDR= 0.05 (Bitton et al., 2015). The red-blue color scale represents the log_2_ fold change expression relative to WT for each given gene and mutant. **C.** Scatterplot of genome-wide log_2_ fold change expression from transcripts in *rrp6*Δ vs *lem2*Δ, both relative to WT. The linear regression line is depicted together with the Pearson correlation coefficient value (*R*). **D.** Expression changes in genes regulated by the nuclear exosome analyzed by RT-qPCR (mostly from the “Mmi1 regulon” (Chen et al., 2011)) in the indicated mutant strains. Additional targets are depicted (HC = heterochromatin controls, EC = euchromatin controls). The color scale represents the log_2_ fold change expression relative to WT for each given gene and mutant. **E.** Clustering based on Pearson’s correlation coefficient (PCC) of RT-qPCR data with genes regulated by the exosome in the indicated mutant strains. **F.** Transcript levels of *sme2*^+^ and *ssm4*^+^ quantified by RT-qPCR in the indicated strains. Data was normalized to *act1*^+^ expression and shown relative to WT on a log_2_ scale. n = 6 independent biological replicates. Individual replicates are shown in a floating bar plot and the line depicts the median. Letters denote different groups from ANOVA and Tukey’s *post hoc* tests at *P* < 0.05.

To confirm that *lem2*Δ and exosome mutants co-regulate transcripts, we performed RT-qPCR with a subset of transcripts, many of them recognized by the meiotic elimination factor Mmi1 (“Mmi1 regulon”, (Chen et al., 2011)) and present in cluster 1 of Fig. 2B. In addition to the aforementioned mutants of exosome factors, we also analyzed mutants lacking the poly(A)-binding protein Pab2 and the MTREC-associated protein Iss10. In agreement with our genome-wide data (Fig. 2B), we found a high level of correlation between *lem2*Δ and *pab2Δ, red1*Δ, and *rrp6*Δ for these selected exosome targets (Fig. 2D,E). Nonetheless, these transcripts were less upregulated in *lem2*Δ than in *rrp6*Δ or *red1*Δ cells (Fig. 2D). We also examined the expression levels of Rrp6 and factors associated with the nuclear exosome. These genes were largely unaltered in *lem2*Δ cells, ruling out that the Lem2-dependent upregulation of exosome targets is indirectly caused by reduced expression of exosome factors (Suppl. Fig. 2F).

We further noticed that genes upregulated in *lem2*Δ, like *mei4*^+^, are often present in Red1-dependent heterochromatin (HC) islands, whereas Red1-independent HC islands were mostly unaffected by *lem2*Δ (with the exception of a ncRNA and an LTR gene; Suppl. Fig. 2G). Red1 is known to associate with chromatin at Red1-dependent HC islands (Zofall et al., 2012), which prompted us to examine the chromatin environment of these loci. While *iss10*^+^ deletion resulted in decreased Red1 binding at the *mei4*^+^ locus, we found no change in *lem2*Δ cells (Suppl. Fig. 2H). Moreover, in contrast to *rrp6*Δ, this locus retained H3K9me2 in the absence of Lem2 (Suppl. Fig. 2I). We also examined H3K9me at heterochromatin domains (HOODs). While deleting *rrp6*^+^ resulted in heterochromatin assembly, as previously reported (Yamanaka et al., 2013), H3K9me2 levels remained low in *lem2*Δ cells (Suppl. Fig. 2I). Together, these findings further support the hypothesis that Lem2 mediates post-transcriptional control of exosome-regulated transcripts, rather than impacting their expression through chromatin changes.

The high resemblance of transcriptional profiles made us wonder whether Lem2 acts through a pathway shared with Pab2, Red1, and Rrp6. To test this, we performed epistasis analysis and examined the expression of prominent exosome targets (*sme2*^+^, *ssm4*^+^, *sno20*^+^, and *snR42*^+^) by RT-qPCR in the single and the respective double mutants. Transcript levels were significantly upregulated in *lem2*Δ compared to WT cells (log_2_ fold-change = 2–5; Fig. 2F, Suppl. Fig. 2J). While these transcripts accumulated to even higher levels in *pab2Δ, red1*Δ and *rrp6*Δ cells (log_2_ fold-change = 4–8), additional deletion of *lem2*^+^ resulted in a non-additive phenotype for *sme2*^+^ and *ssm4*^+^, suggesting an epistatic interaction (Fig. 2F). This was different for sno/snRNAs, for which we found an additive increase in *lem2*Δ *pab2*Δ and *lem2*Δ *red1*Δ double mutants (Suppl. Fig. 2J). This further supports the idea that Lem2 plays a broad role in RNA degradation by collaborating with distinct pathways with different substrate specificity.

### Lem2 interacts with exosome-targeting factor Red1

Given the epistatic interaction between Lem2 and Red1 (Fig. 2F), we wondered whether they directly interact. To address this question, we generated strains expressing Lem2-GFP and Red1-6xHA expressed from their endogenous loci analogous to previous studies (Barrales et al., 2016; Shichino et al., 2020). Using co-immunoprecipitation (coIP) experiments, we found that Lem2 binds Red1 *in vivo* (Fig. 3A). Similar to MTREC interactions with other factors (Shichino et al., 2020; Zhou et al., 2015), this physical association was insensitive to RNase and Benzonase treatment, suggesting that the interaction is not mediated by RNA or DNA (Fig. 3A).

**Fig. 3:**
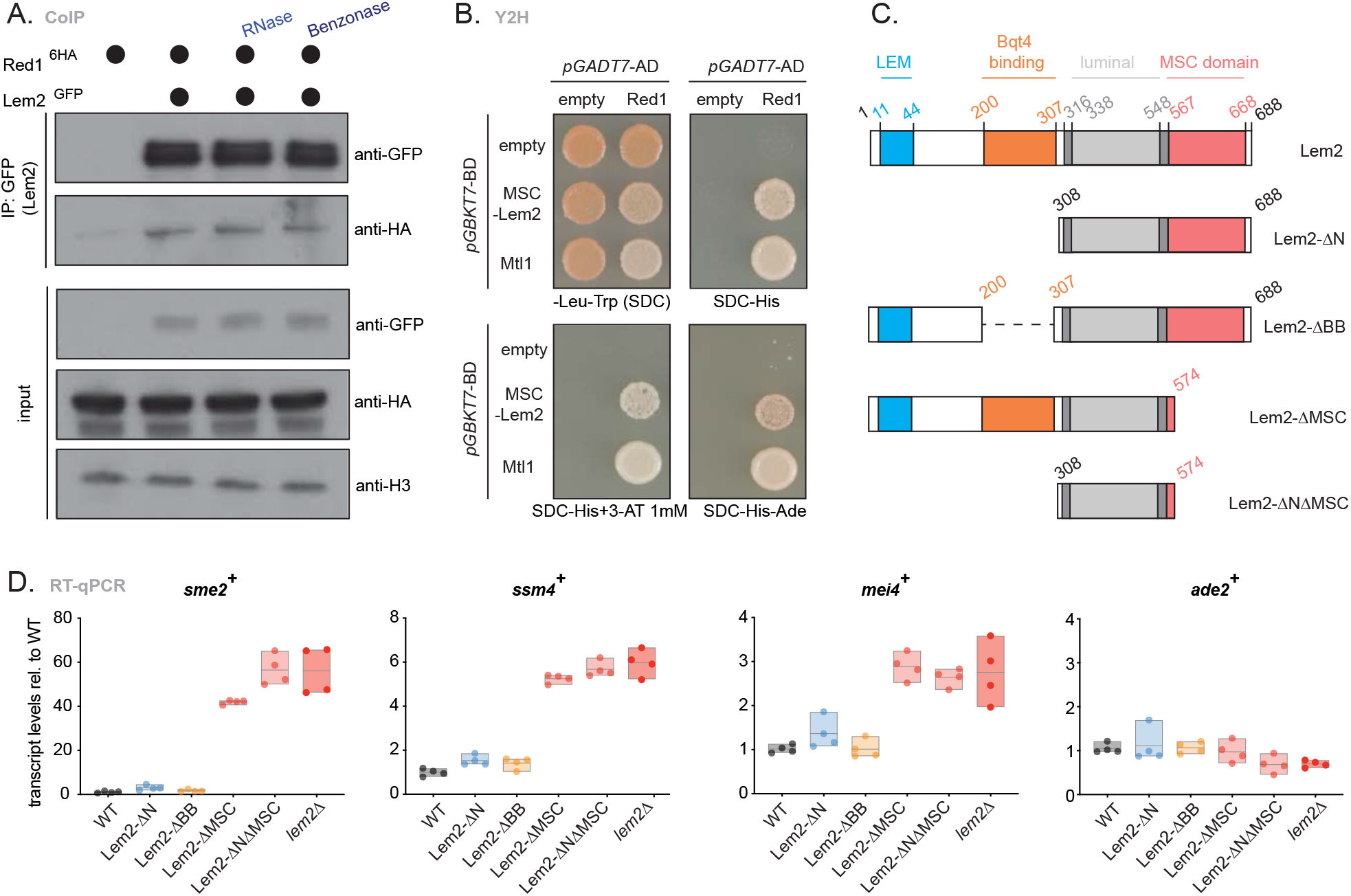
Lem2 physically interacts with the nuclear exosome through the MSC domain. **A.** Co-immunoprecipitation of Red1-6xHA with Lem2-GFP in untreated cells, or cells treated with RNase or benzonase. H3 served as loading control. **B.** Y2H analysis of Red1 with MSC-Lem2 or Mtl1, grown for 3 days on medium with increasing stringency (SDC, SDC-His, SDC-His + 3-AT 1 mM, SDC-His-Ade). Fusions with Gal4 activation domain (*pGADT7-AD*) or Gal4 DNA-binding domain (*pGBKT7-BD*) are shown. **C.** Schematic representation of Lem2 truncation constructs. Protein domains are highlighted along with their amino acid length. All these constructs were C-terminally GFP-tagged. **D.** Transcript levels of *sme2^+^, ssm4^+^, mei4*^+^ and *ade2*^+^ quantified by RT-qPCR on the Lem2 truncation mutants shown in Fig. 3c. Data normalized to *act1*^+^ expression is shown relative to WT. n = 4 independent biological replicates. The individual replicates are shown in a floating bar plot and the line depicts the median.

Lem2 has distinct structural domains that mediate different functions. The N-terminal LEM domain contributes to centromere association and clustering at the NE, whereas the C-terminal MSC domain is required for telomere anchoring and heterochromatin silencing (Barrales et al., 2016). In addition, a small region located adjacent to the first transmembrane domain was shown to interact with the integral membrane protein Bqt4 (Hirano et al., 2018; Hu et al., 2019). We assessed the role of the N- and C-terminal Lem2 regions in Red1 binding using truncated mutants (Lem2-ΔN and Lem2-ΔMSC, respectively; Fig. 3C). The absence of the Lem2 N-terminus did not impair Red1 association (Suppl. Fig. 3A). In contrast, removal of the C-terminal MSC domain abolished Red1 binding, indicating that it is essential for Lem2 interaction with Red1 (Suppl. Fig. 3A). To test whether the MSC domain is sufficient for Red1 association, we performed yeast two-hybrid (Y2H) assays using the MSC domain of Lem2 and full-length Red1. Consistent with our coIP results, we observed a robust interaction between Lem2-MSC and Red1 (Fig. 3B), arguing that the MSC domain is necessary and sufficient to mediate Red1 interaction, since the MTREC complex is not present in budding yeast. We further observed association of Lem2-MSC with Iss10, but not with Rrp6, Pab2 or Mtl1 (Suppl. Fig. 3B). Together, these results suggest that Lem2 directly mediates the interaction with MTREC.

To examine whether the domain within Lem2 critical for Red1 interaction is also crucial for exosome substrate repression, we analyzed transcript levels of selected exosome targets by RT-qPCR using a series of previously described Lem2 truncation mutants ((Hirano et al., 2018); Fig. 3C). Whereas truncations in the N-terminal region of Lem2 did not affect expression levels, mutants lacking the C-terminal region largely phenocopied the silencing defect of *lem2*Δ cells (Fig. 3D). To further test whether the perinuclear localization of the MSC domain is important for the repression of exosome targets, we generated a soluble Lem2 protein comprising the MSC domain without transmembrane domains and fused to GFP and the SV40 nuclear localization signal (NLS) (Suppl. Fig. 3C). Similar to our previous study using overexpressed MSC-GFP (Barrales et al., 2016), endogenously expressed MSC-GFP produced a diffuse nuclear pattern, which differs from the rim shape of full-length Lem2 (Suppl. Fig 3D). The soluble MSC domain was expressed at lower levels than full-length Lem2, but was nonetheless detectable (Suppl. Fig. 3D,E). Analogous to heterochromatin transcripts (Barrales et al., 2016), the soluble MSC domain failed to suppress the accumulation of meiotic transcripts in *lem2*Δ cells (Suppl. Fig. 3F), indicating that proper NE location of the MSC domain is crucial for regulation of exosome target RNAs. Based on these results, we conclude that Lem2 cooperates with the nuclear exosome by directly interacting with Red1 *in vivo* through its MSC domain at the nuclear periphery.

### Lem2 regulates silencing of exosome targets at the nuclear periphery

Our data (Suppl. Fig. 3F) and previous studies (Barrales et al., 2016; Tange et al., 2016) indicate that perinuclear location of Lem2 is critical for its gene repression function. This implies that targeting or degradation of Lem2-dependent RNA substrates by the nuclear exosome takes place at the nuclear periphery. To test this hypothesis, we used an engineered strain that expresses a reporter mRNA containing 14 DSR copies and 4 copies of the U1A snRNA stem loop (Shichino et al., 2018). This DSR reporter can be visualized by co-expression of U1A-YFP, resulting in a single dot of the DSR transcript (Fig. 4A,C; (Shichino et al., 2018)). Using RT-qPCR, we confirmed that the presence of DSR sequences mediates reporter transcript elimination in an Iss10- and Red1-dependent manner (Suppl. Fig. 4A). Lack of Lem2 also resulted in a moderate but reproducible upregulation of the DSR transcript levels (Fig. 4B), verifying that Lem2 regulates the expression of this synthetic exosome substrate.

**Fig. 4:**
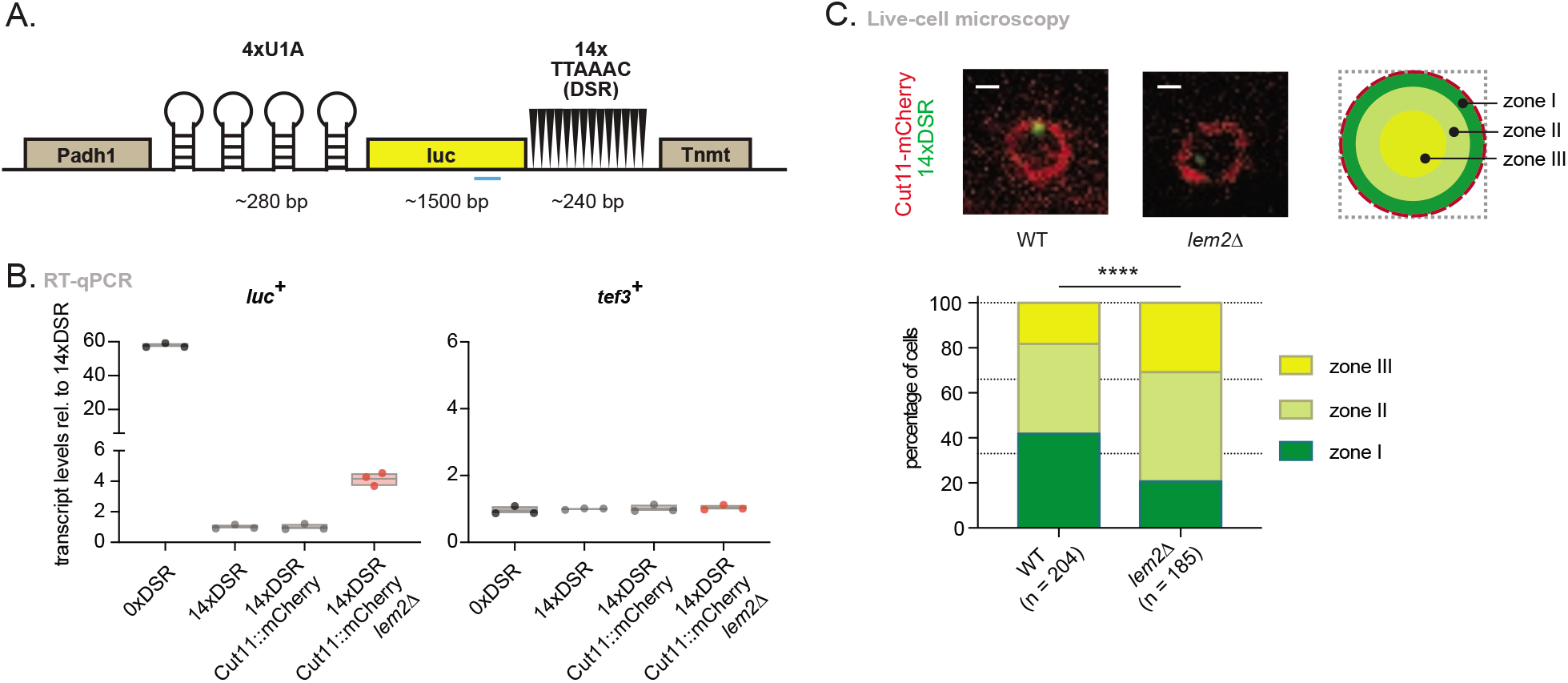
Exosome substrates localize at the nuclear periphery. **A.** Schematic representation of the engineered DSR containing construct, adapted from (Shichino et at., 2018). 14 copies of DSR are expressed together with 4 copies of the U1A tag and the luciferase ORF and are regulated by the *adh1* promoter (Padh1) and the *nmt1* terminator (Tnmt). **B.** Transcript levels of *luc*^+^ and *tef3*^+^ quantified by RT-qPCR on a strain encoding 0xDSR copies (0xDSR), 14xDSR copies (14xDSR), 14xDSR and the NE marker in a WT (14xDSR Cut11::mCherry) or *lem2*Δ background (14xDSR Cut11::mCherry *lem2*Δ). Data was normalized to *act1*^+^ expression and shown relative to the 14xDSR strain. n = 3 independent biological replicates. The individual replicates are shown in a floating bar plot and the line depicts the median. **C.** Top left: Live-cell microscopy representative images of the DSR containing strain (Fig. 4a) in a WT or *lem2*Δ background. Cut11-mCherry marks the NE. A single z-stack is shown. Top right: schematic representation of the division of a *S. pombe* nucleus in three areas with equal surfaces. These zones are called I-III depending on how close to the periphery they are. Bottom: quantification of DSR location in WT and *lem2*Δ backgrounds relative to the periphery expressed in percentage of cells. n = number of cells counted in two independent experiments. **** = P < 0.0001 from χ^2^ test analysis. Scale bar = 1 μm.

Using live-cell imaging, we next studied the localization of the DSR-containing RNA reporter relative to the nuclear periphery, which was marked with Cut11-mCherry (Suppl. Fig. 4B) in WT and *lem2*Δ strains. We determined the frequency at which the DSR dot localizes to specific nuclear areas using a zoning assay (Hediger et al., 2004). Interestingly, the DSR dot showed preferential localization to the nuclear periphery in about 40% of WT cells (zone I). This pattern, however, was significantly perturbed in the *lem2*Δ mutant, displaying perinuclear localization in only about 20% of the cells, indicating that Lem2 promotes the perinuclear localization of this exosome RNA substrate (Fig. 4C). This differed from the localization of genomic loci, as shown for the lacO array/GFP-lacI marked *sme2*^+^ locus, which did not appear to be preferentially localized at the periphery, (Suppl. Fig. 4C). These data indicate that regulation of exosome targets by Lem2 takes place at the nuclear periphery.

### Lem2 assists exosome substrate targeting

Since Lem2 contributes to the repression and localization of exosome RNA substrates (Fig. 2 and Fig. 4) and physically interacts with Red1 (Fig. 3), we speculated that Lem2 might assist in the degradation process through transcript recruitment and handover to MTREC for processing. While the N-terminal region of Lem2 associates with centromeric chromatin (Banday et al., 2016; Barrales et al., 2016; Tange et al., 2016), the MSC domain of the human Lem2 homolog MAN1 (hMAN1) has been shown to bind DNA *in vitro* (Caputo et al., 2006). Furthermore, hMAN1 also contains an RNA recognition motif as a C-terminal extension (Brachner et al., 2005). However, whether Lem2 can bind RNA is unknown.

We performed RNA immunoprecipitation (RIP) assays followed by RT-qPCR to examine the potential association of Lem2 with exosome substrates. We assessed binding of *sme2*^+^ or *ssm4*^+^ transcripts, which contain several DSR sequences and are recognized by members of the nuclear exosome pathway (Hiriart et al., 2012) (Fig. 5A). However, we were unable to detect those transcripts bound to the immunoprecipitated Lem2-GFP (Fig. 5B). We therefore tested the alternative hypothesis that Lem2 plays an accessory role in substrate recognition or loading RNAs onto exosome-targeting factors. To this end, we expressed GFP-Mmi1 and Red1-myc from their endogenous loci and confirmed that the presence of epitope tags does not interfere with their function (Suppl. Fig. 5). Using RIP, we found that *sme2*^+^ and *ssm4*^+^ transcripts were abundantly enriched with Mmi1 and Red1 (20-60 fold and 150-250 fold, respectively. See Fig. 5C,D), in agreement with previous reports (Harigaya et al., 2006; Sugiyama and Sugioka-Sugiyama, 2011). Strikingly, deletion of *lem2*^+^ markedly reduced or abolished binding of these transcripts to both Mmi1 and Red1 (Fig. 5C,D), revealing that Lem2 plays a critical role in the early step of RNA recognition.

**Fig. 5:**
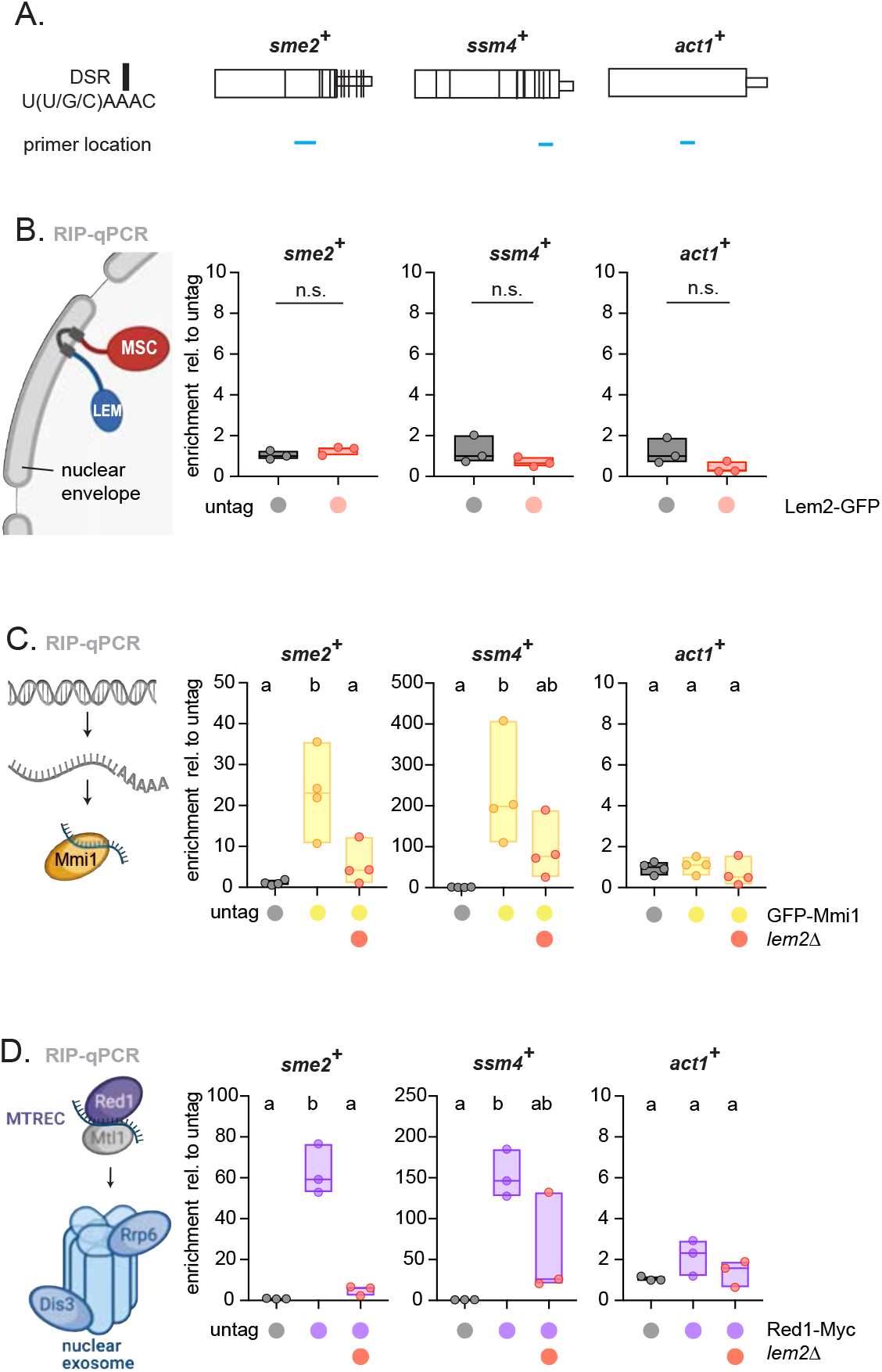
Lem2 promotes binding of RNA targets with exosome-targeting factors. **A.** Schematic representation of DSR containing genes *sme2*^+^, *ssm4*^+^ and control gene actin (*act1*^+^). Sizes were made relative to match lengths. DSR location was adapted from (Chen et al., 2011). DSR location is indicated with black lines over the genes. qPCR primer location is indicated by blue lines below the genes. **B.** Left: Scheme of Lem2 at the NE depicting its functional LEM and MSC domains. Right: transcript binding to Lem2-GFP as assayed by RIP-qPCR analysis. T-test analysis was performed. **C, D.** Left: schematic representations of exosome machinery components Mmi1 and Red1. Right: transcript binding to GFP-Mmi1 (C) or Red1-Myc (D) analyzed by RIP-qPCR analysis in WT and *lem2*Δ cells. For B, C, D, the data was divided by the input and is shown relative to the median of the untagged strain. n = 3-4 independent biological replicates. The individual replicates are shown in a floating bar plot and the line depicts the median. For C and D letters denote different groups from ANOVA and Tukey’s post hoc tests at P < 0.05.

### Multiple pathways contribute to exosome-mediated RNA degradation

Various exosome factors, including Mmi1, Red1 and Rrp6, assemble during vegetative growth into single or multiple nuclear foci, which have been proposed to act as sites of RNA degradation (Shichino et al., 2020; Sugiyama and Sugioka-Sugiyama, 2011; Sugiyama et al., 2016; Yamashita et al., 2013). While Mmi1-containing EMC and MTREC can form independent foci, their mutual interaction depends on Iss10, which physically interacts with Red1 (Yamashita et al., 2013; Zhou et al., 2015). We investigated whether the formation and localization of these nuclear foci requires Lem2. Using livecell imaging, we confirmed that the exosome-targeting factors Red1, Erh1 and Mmi1 form nuclear foci. However, we observed neither a perinuclear enrichment of these foci in WT cells nor a change in their formation or localization upon deletion of *lem2*^+^ (Suppl. Fig. 6A). We further examined whether Lem2 is critical for the interaction between Red1 and Mmi1, analogous to the role of Iss10 in bridging Mmi1 recruitment to Red1 (Lee et al., 2013; Sugiyama and Sugioka-Sugiyama, 2011; Yamashita et al., 2013; Zhou et al., 2015). As previously reported (Shichino et al., 2020; Sugiyama and Sugioka-Sugiyama, 2011), we found that GFP-Mmi1 coimmunoprecipitates with Red1-6xHA; however, association was unaffected by *lem2*^+^ deletion (Suppl. Fig. 6B). Since *erh1*Δ and *iss10*Δ mutants also showed little functional overlap with *lem2*Δ (Fig. 2), we tested the hypothesis that RNA degradation involving Lem2 is mediated through an Erh1- and Iss10-independent pathway. When examining transcript levels in the *ehr1Δ lem2*Δ double mutant, we found a synthetic increase for several meiotic transcripts but not for snoRNAs, which are processed independently of the Mmi1 elimination pathway (Fig. 6A). Lem2 also acts redundantly with Iss10 in eliminating *mei4*^+^ and *ssm4*^+^ transcripts (Fig. 6B). Interestingly, this was different for *sme2*^+^ and *mei3*^+^ transcripts, which were exclusively controlled by Lem2 (Fig. 6B). We further observed additive phenotypes for *lem2*Δ in combination with *air1*Δ (defective in TRAMP-mediated degradation) (Suppl. Fig. 6C). Since stabilization of *meiRNA* and its assembly into the Mei2 dot during early meiosis inactivates Mmi1-dependent elimination (Harigaya and Yamamoto, 2007), we tested whether other exosome targets accumulating in *lem2*Δ cells depend on Mei2 dot formation. However, additional deletion of *sme2*^+^ did not suppress the accumulation of meiotic transcripts or snoRNAs in *lem2*Δ cells (Suppl. Fig. 6D), suggesting a direct role of Lem2 in the degradation of these exosome targets. Together, these results imply that RNA degradation is coordinated through distinct degradation pathways at the nuclear periphery and nuclear bodies, which differ in their substrate specificity and depend on Lem2 and Iss10, respectively.

**Fig. 6:**
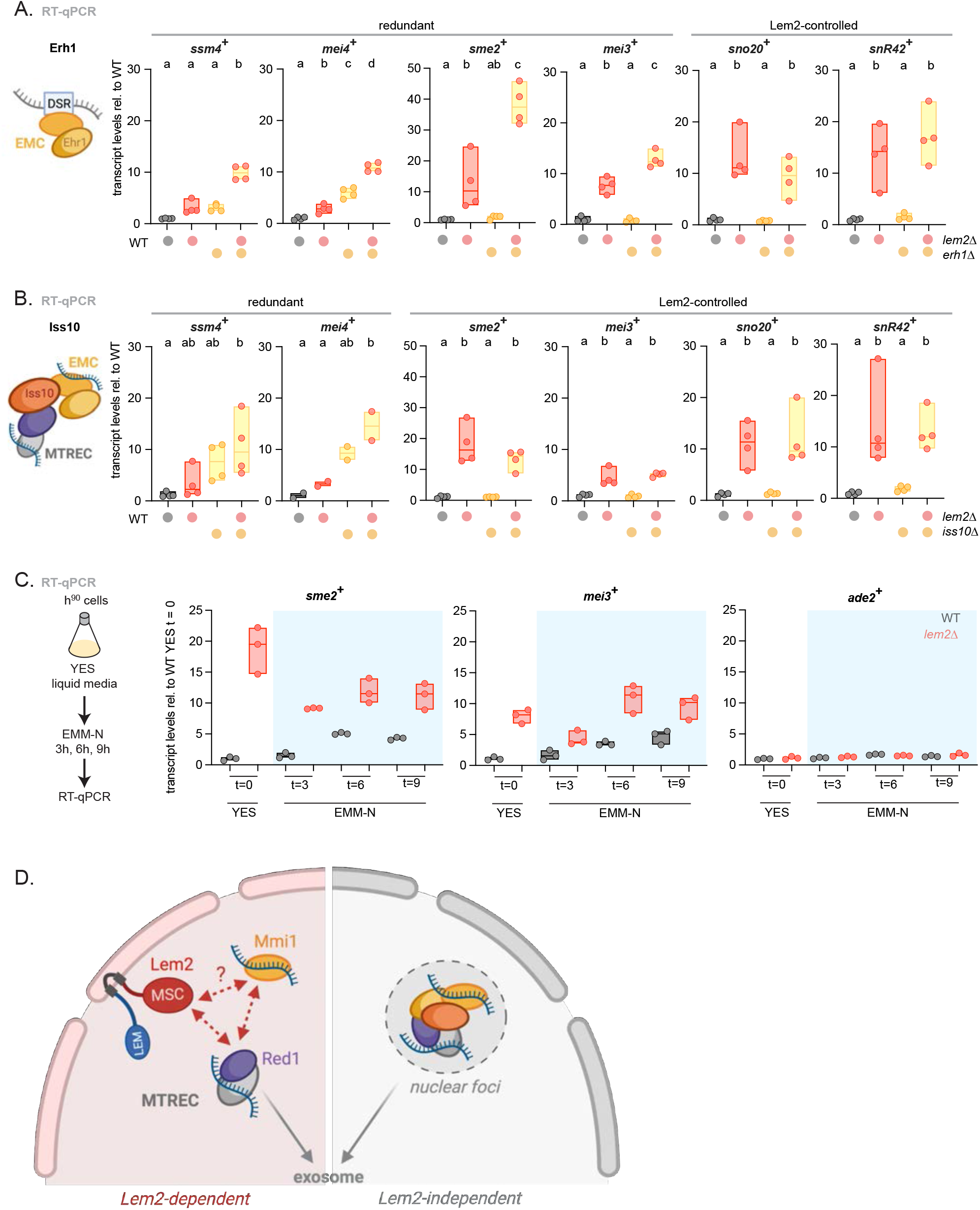
RNA regulation by Lem2 at the nuclear periphery occurs independently of exosome factors associated with nuclear foci. **A, B.** Left: scheme of the subunit that was mutated on its own or together with Lem2. Right: transcript levels of *ssm4^+^, mei4^+^, sme2^+^, mei3^+^, sno20*^+^ and *snR42*^+^ quantified by RT-qPCR in the indicated strains (A = *erh1*Δ, B = *iss10*Δ). Redundant or Lem2-controlled substrates are indicated above. **C.** Left: scheme showing the experimental setup and media shift. Right: transcript levels of *sme2^+^, mei3*^+^ and *ade2*^+^ quantified by RT-qPCR in the indicated strains (grey = WT, red = *lem2*Δ), time points (t = 0, 3, 6 or 9 hours) and media (YES or EMM-N). Data was normalized to euchromatin expression and shown relative to the WT strain at t = 0 in YES. n = 3 independent biological replicates. **D.** Model. Left: Lem2 cooperates with the nuclear exosome machinery at the nuclear periphery by directly interacting with Red1 and assisting in substrate binding by Mmi1 and MTREC. Absence or inactivation of Lem2 results in impaired DSR substrate location at the periphery, reduced binding to exosome-targeting factors, and eventually an increase in untimely transcript expression due to lower RNA degradation (Lem2-dependent). Lem2 may also interact with other factors associated with Mmi1. Right: Nuclear foci are formed by multiple exosome factors in an Iss10-dependent manner and likely represent additional sites of RNA degradation (Lem2-independent). Both pathways show partial redundancy for most meiotic transcripts, while ncRNAs are mainly degraded by the Lem2-dependent pathway. For A, B and C, the individual replicates are shown in a floating bar plot and the line depicts the median. For A and B data was normalized to *act1*^+^ expression and shown relative to the WT strain. n = 2-4 independent biological replicates. Letters denote different groups from ANOVA and Tukey’s *post hoc* tests at *P* < 0.05.

Nitrogen starvation promotes sexual differentiation and entry into meiosis. Accumulation of meiotic transcripts coincides with Iss10 degradation and disassembly of nuclear foci (Wei et al., 2021). We therefore speculated that Lem2-dependent degradation of meiotic transcripts might also be controlled during meiotic onset. Previously, Lem2-dependent heterochromatin assembly and minichromosome stability were reported to be regulated by nutrient availability (Tange et al., 2016). In particular, when cells are grown in minimal medium (Edinburgh minimal medium, EMM), Lem2 association with centromeres is abolished (Tange et al., 2016). Furthermore, heterochromatic transcripts are no longer repressed in a Lem2-dependent manner in EMM (Suppl. Fig. 6E), suggesting that Lem2 becomes inactivated under such growth conditions. This situation is reminiscent of the switch from mitotic to meiotic growth, as EMM also induces meiosis in post-logarithmic cell cultures (Simonetti et al., 2017). To investigate whether Lem2 activity is regulated during meiotic onset, we grew mating-competent h^90^ cells in rich medium and then analyzed *sme2*^+^ and *mei3*^+^ transcripts upon transfer to EMM lacking nitrogen in a time-course experiment (Fig. 6C). As predicted, we observed a steady increase of these meiotic transcripts in WT cells upon nitrogen starvation, reaching a plateau after 6 hours (Fig. 6C). In contrast, meiotic transcripts were already upregulated in *lem2*Δ cells in rich medium and did not further increase when starved (Fig. 6C). The non-additive phenotype suggests that Lem2 contributes to meiotic transcript accumulation in WT cells. While we noticed that these meiotic transcripts did not accumulate to the same level in WT as in *lem2*Δ cells, we cannot exclude that some cells within the population might not have entered the meiotic program, likely explaining the weaker phenotype in WT cells. Based on these results, we propose that nutrient-dependent inactivation of Lem2 functionally contributes to the fine-tuning of meiotic transcript accumulation during early meiosis. Beyond this role in meiosis, Lem2 is key to a spatially and functionally distinct pathway that operates in post-transcriptional regulation at the NE.

## Discussion

Nuclear architecture plays a key role in transcriptional regulation. Transcriptionally silent chromatin often localizes to the nuclear periphery, which is thought to provide a specialized compartment for gene repression (Akhtar and Gasser, 2007). However, whether the NE is also important for broader modes of gene regulation is largely unknown. In this study, we demonstrate that the conserved INM protein Lem2 collaborates with the nuclear exosome to repress meiotic transcripts and ncRNAs. Several lines of evidence suggest that this novel role of Lem2 takes place at the post-transcriptional level and is distinct from its previously described function in heterochromatin silencing (Barrales et al., 2016; Tange et al., 2016): (1) Whereas Lem2 cooperates with silencing factors (Clr4/HP1, RNAi, SHREC) at heterochromatic loci, those factors have little impact on the repression of targets co-regulated by Lem2 and the exosome. (2) While *lem2*^+^ deletion causes accumulation of various targets (e.g., the ncRNA *sme2*^+^ and the heterochromatic transcript *tlh1*^+^), Pol II abundance is increased at heterochromatin but not at ncRNA genes, separating the two functions of Lem2. (3) Mutants lacking both Lem2 and exosome factors (Rrp6, MTREC) display non-additive phenotypes for the repression of several meiotic transcripts. (4) Lem2 physically interacts with the MTREC subunit Red1 via its MSC domain *in vivo*. (5) Lem2 affects the perinuclear localization of an RNA substrate but not the genomic *sme2*^+^ locus. (6) Lem2 promotes the binding of RNA substrates to exosome targeting complexes (EMC, MTREC). Based on these findings, we propose that Lem2 recruits exosome co-factors to the nuclear periphery to coordinate post-transcriptional RNA processing and degradation, in addition to its role in recruiting heterochromatin factors for transcriptional silencing.

Despite major differences in transcriptional and post-transcriptional regulation, from a mechanistic perspective, Lem2’s role in RNA degradation is reminiscent of its function in heterochromatin silencing by increasing the local concentration of repressors at the nuclear periphery. As previously shown, Lem2 promotes the recruitment of the repressor complex SHREC to heterochromatin (Barrales et al., 2016). Similarly, we find that Lem2 facilitates the association of MTREC with RNA substrates (Fig. 5) and interacts with Red1 through its MSC domain (Fig. 3). The association between Lem2 and Red1 both in the presence of nucleases or when heterologously expressed in budding yeast implies a direct interaction (Fig. 3). We further find that Lem2 interacts via its MSC domain with another MTREC component, Iss10, although the physiological relevance of this interaction remains to be determined (see below). Together, this suggests that Lem2 employs a common mechanism for interaction with its downstream partners, which is in agreement with the MSC-dependent recruitment of members of the ESCRT pathways in NE repair and other functions reported for Lem2 homologs (Appen et al., 2020; Capella et al., 2020; Gu et al., 2017; Pieper et al., 2020; Thaller et al., 2019). Thus, Lem2 molecules may create a general recruitment platform at the INM for interaction with various partners. How Lem2 specifically coordinates these different functions remains unknown. Interestingly, while heterochromatin silencing and RNA degradation are mediated specifically by Lem2, but not its paralog Man1 *in vivo* ((Barrales et al., 2016); this study), we found that the MSC domain of Man1 is still capable of interacting with Red1 in Y2H assays (Suppl. Fig. 3G). This implies that the MSC domains of Lem2 and Man1 are not sufficient to discriminate between different binding partners and that additional factors are likely involved in regulating the Lem2 association with effector proteins *in vivo*.

Various membrane-less structures, known as nuclear bodies, have been described and associated with specific functions, such as the Cajal body involved in snRNA and snoRNA modification and assembly (Mao et al., 2011). RNA turnover has also been proposed to occur within subnuclear foci into which exosome factors assemble in an Iss10-dependent manner (Shichino et al., 2020; Sugiyama and Sugioka-Sugiyama, 2011; Sugiyama et al., 2016; Yamashita et al., 2013). This raises the question of how RNA degradation in these nuclear foci relates to Lem2-mediated regulation at the NE. Although we found that an exosome substrate localizes more frequently at the nuclear periphery in WT than in *lem2*Δ cells (Fig. 4), such localization was not seen for Red1, Erh1, or Mmi1 foci (Suppl. Fig. 6). This was surprising, as this engineered substrate was previously reported to co-localize with Mmi1 (Shichino et al., 2018). However, this reporter construct largely forms a single focus in the nucleus (our data and (Shichino et al., 2018)), whereas Mmi1 assembles into multiple foci. Similarly, Mmi1 forms a distinct focus with the non-coding *mamRNA* that differs from other Mmi1 foci (Andrić et al., 2021), implying that Mmi1 assembles into different nuclear bodies with distinct functions. With the employed method, we cannot distinguish whether individual foci display different distributions within the nucleus; thus, it might be possible that individual Mmi1 foci co-localizing with the reporter substrate preferentially localize at the periphery. Nonetheless, we surmise that the majority of EMC and Red1 foci do not co-localize with the NE and may represent independent degradation pathways that differ from the regulation by Lem2. Indeed, *lem2*Δ mutants additionally lacking Erh1 or Iss10 display a synergistic upregulation of meiotic transcripts (Fig. 6). This suggests that multiple RNA degradation pathways act in parallel. This likely explains why the phenotype of *lem2*Δ is weaker than that of *red1*Δ and *rrp6*Δ mutants (see model, Fig. 6D). The idea of having redundant pathways would further solve the apparent discrepancy reported for Iss10 being critical for nuclear foci formation while only playing a minor role in the elimination of meiotic transcripts (Egan et al., 2014; Shichino et al., 2020; Wei et al., 2021; Yamashita et al., 2013). We speculate that separate pathways allow differential regulation and distinct substrate specificity. Indeed, degradation of the meiotic transcripts *mei3*^+^ and *sme2*^+^ is predominantly controlled by Lem2 but not Iss10. Conversely, based on their physical interaction (Suppl. Fig. 3), Lem2 and Iss10 might also cooperate in the degradation of other substrates. We further observe that Lem2 functions in part independent of MTREC in the degradation of sno/snRNAs (Suppl. Fig. 2), indicating that additional Lem2-dependent pathways exist. Further work is needed to elucidate Lem2’s specific role in the broad spectrum of its substrates.

The expression of meiotic genes is toxic for cells during vegetative growth and therefore tightly regulated (Harigaya and Yamamoto, 2007; Wei et al., 2021). Under nutrient starvation, cells mate and undergo meiosis, which requires the timely orchestration of the meiotic gene expression program (Mata et al., 2002). Nitrogen starvation is signaled by inactivation of the TOR pathway, resulting in the dephosphorylation and degradation of Iss10 by the ubiquitin-proteasome system (Wei et al., 2021). Similarly, cells shifted from rich to minimal growth medium no longer show Lem2-dependent transcriptional silencing (Suppl. Fig. 6E). Likewise, WT cells undergoing meiosis accumulate meiotic transcripts to almost the same levels as *lem2*Δ cells in minimal medium, suggesting that the upregulation of these transcripts can, at least in part, be attributed to Lem2 inactivation (Fig. 6C). Hence, we propose that Lem2 is part of a regulatory circuit that fine-tunes gene expression in response to environmental cues. Interestingly, a similar nutrient-dependent regulation is seen for Lem2 function in heterochromatin maintenance (Tange et al., 2016). Of note, nitrogen depletion causes a dramatic change in chromatin organization, with some loci losing their peripheral localization and moving towards the interior of the nucleus (Alfredsson-Timmins et al., 2009). However, we noted that Lem2 inactivation is triggered in minimal medium even in the presence of nitrogen, suggesting that a signaling cascade other than the TOR pathway is involved. How different growth conditions alter Lem2 activity yet remains unknown. Since neither transcript levels (Suppl. Fig. 6F) nor protein levels of Lem2 (Tange et al., 2016) are altered under minimal growth conditions, Lem2 might be subject to post-translational modifications. Such modifications could affect its location within the NE or its association with downstream factors. Alternatively, minimal growth conditions may alter the expression of Lem2’s downstream partners. Thus, it will be interesting to address how nutritional cues affect Lem2 function and elucidate the underlying signaling cascade.

Given that both Lem2 and exosome-targeting complexes are found in higher eukaryotes, we propose that this pathway is broadly conserved. Indeed, the nuclear exosome has been shown to localize to the NE in other organisms, such as *Drosophila* (Graham et al., 2006). Further studies examining the nuclear location of the exosome in fission yeast and higher eukaryotes may shed light on the collaborating mechanisms that help regulate this complex machinery. However, our data demonstrate that RNA degradation is not a generic process, but a spatially specific mode of regulation that is critical in the biological response to environmental change.

## Methods

### Yeast techniques, plasmids and strains

A list of the strains used in this study can be found in Suppl. Tables 3 and 4.

Standard media and genome engineering methods were used. Cells were grown in rich medium (YE5S aka YES) except for the data shown in Fig. 3D, 6A, Supp. Fig. 6E,F (cells were initially grown in minimal medium (EMM) then shifted for 12 h into YES) and Fig. 6C (cells were initially grown in YES then shifted into EMM-N). Strains expressing constructs derived from *pREP81* vectors (shown in Suppl. Fig. 3A) were grown in EMM-leu.

Endogenously tagged strains were generated through homologous recombination using *pFA6a* or *pYM*-based vectors (Janke et al., 2004). For the endogenously tagged Lem2-GFP strains, the respective fragments were cloned into a *pJK210* vector and integrated into *lem2*Δ background strains (Suppl. Table 5).

### RNAseq library preparation and data analysis

For RNAseq, 1 μg of RNA was used as starting material to make libraries following manufacturer instructions for NEBNext Ultra Directional RNA Library Prep Kit for Illumina (NEB). Two or more biological replicates were used for making libraries in parallel. For *man1*Δ *and clr4*Δ, three biological replicates were processed. Single-end, 50 bp sequencing of libraries was performed on a HiSeq1500 sequencer in the LAFUGA core facility, in the Gene Center, Munich. Raw reads were de-multiplexed using Je (v1.2). Adapter-trimmed reads were aligned to the *S. pombe* reference genome (ASM294v2.27) and a custom GTF file using STAR (v2.7.3a), then processed using RSEM (v1.3.3). Differential expression was analyzed using the DESeq2 (v1.22.2) and tximport (v1.10.1) R libraries. For PCA plotting, data was batch normalized using the sva (v3.30.1) R library. Bedgraph coverage files for plus and minus strands were generated using genomecov (bedtools v2.29.1).

### RT-qPCR analyses

RT-qPCR experiments were performed as previously described (Braun et al., 2011). Data from 2-6 independent biological replicates are shown as individual data points with a line depicting the median. cDNAs were quantified by qPCR using the prima-QUANT CYBR Master Mix (Steinbrenner Laborsysteme) in a QuantStudio 5 or 3 Real-Time PCR System (Applied Biosystems). The primers used are listed in Suppl. Table 6. Day normalization was performed as previously described (Georgescu et al., 2020). When using double mutants, statistical testing was performed using R. Multiple testing was performed using ANOVA and Tukey’s *post hoc* test at a 0.05 significance level.

### Y2H assays

The specified constructs were cloned into either *pGADT7* or *pGBKT7* vectors (Clontech™). The *Saccharomyces cerevisiae* strain Y2H Gold (Takara) was used to co-transform the plasmids following manufacturer’s instructions. Spotting assays were performed 3-5 days after transformation of the plasmids. Different dropout mixes were used to assess the strength of the interaction: SDC-Leu-Trp (Formedium), SD-Leu-Trp-His (Formedium), SD-Leu-Trp-His+1mM 3-AT (3-aminotriazol, Sigma) and SD-Leu-Trp-His-Ade (Formedium).

### Co-immunoprecipitation assays

CoIP assays were performed as previously described (Capella et al., 2020) with a few modifications in the lysis buffer, as follows. The lysis buffer contained 50 mM HEPES pH 7.4, 100 mM NaCl, 10% glycerol, 1 mM EDTA pH 8, 2.5 mM MgCl_2_, 0.5% NP-40, 1x complete EDTA-free protease inhibitor cocktail (Roche), 2 mM PMSF (Serva), 20mM N-ethylmaleimide (NEM, Sigma). Immunoprecipitation was performed using GFP-trap or Myc-trap (Chromotek). While immunoprecipitation took place, RNase (Roche) or benzonase (Sigma) was added in the specified assays. Beads were then washed four times with lysis buffer and two times with wash buffer (50 mM Tris pH 7.5, 100 mM NaCl, 1 mM EDTA pH 8, 2.5 mM MgCl_2_). Proteins were eluted and analyzed by immunoblotting.

### Immunoblotting

Cells corresponding to OD_600_ = 1 (≈ 2 x 10^7^ cells) were pelleted from a suspension culture grown to mid-log phase. Total protein extracts were made using trichloroacetic acid (TCA, Sigma) precipitation (Knop et al., 1999). Proteins were solubilized in HU buffer (8 M urea, 5% SDS, 200 mM Tris-HCl pH 6.8, 20 mM dithiothreitol (DTT, Sigma) and bromophenol blue 1.5 mM). Proteins were resolved on NuPAGE 4%-12% gradient gels (Invitrogen) or self-made 8% gels. Proteins were transferred onto PVDF membranes (polyvinylidene fluoride membranes, GE-Healthcare) and analyzed by standard immunoblotting techniques using specific antibodies. H3 antibody was used as loading control

### Antibodies

Monoclonal anti-HA antibody (3F10, 1:1000) was purchased from Roche. Monoclonal anti-GFP (B-2, 1:1000) was purchased from Santa Cruz Biotechnology. Monoclonal anti-H3 (1B1-B2, 1:5000) was purchased from Active Motif. Polyclonal anti-Myc (ab9106, 1:2000) antibody was kindly provided by M. Spletter, purchased from Abcam. Secondary antibodies fused to HRP were used for detection (Goat anti-mouse HRP 1:3000, BioRad; Goat anti-rat HRP 1:3000, Merck Millipore; Goat anti-rabbit HRP 1:3000, BioRad)

### Live-cell microscopy

Live cell imaging was essentially performed as described (Barrales et al., 2016). In brief, cells were grown overnight on rich medium to logarithmic phase. Prior to imaging, cells were attached with lectin (Sigma) to glass bottom dishes with a micro well (MatTek). Cells were imaged on a Zeiss AxioObserver Z1 confocal spinning disk microscope with an EMM-CCD camera (Photometrics, Evolve 512) through a Zeiss Alpha Plan/Apo 100x/1.46 oil DIC M27 objective lens. Z-stacks were obtained at focus intervals of 0.4 μm. Fiji/ ImageJ software was used to measure the distances between the foci and the periphery.

### RIP assays

Cell lysates prepared from equal amounts of cells (between 100-165 OD600) were fixed with 1% formaldehyde (Sigma) for 15 min at RT, followed by quenching 5 min at RT with 125 mM glycine (Sigma). Cultures were spun down, washed once with 1xPBS and frozen in liquid nitrogen. Cells pellets were resuspended in lysis buffer (250 mM KCl, 1% Triton X-100, 0.1% SDS, 0.1% Na-deoxycholate, 50 mM HEPES pH 7.5, 2 mM EDTA, 2 mM EGTA, 5 mM MgCl_2_, 0.1% Nonidet P-40, 20% glycerol) and lysed with glass beads (Roth) in a bead beater (Precellys 24, Peqlab). Fragmented material was sonicated (Qsonica Q800R1) for 1h with cycles of 30s ON/OFF at 4°C. The lysate was cleared and used for immunoprecipitation with 15 μl GFP-trap or Myc-trap (Chromotek). Beads were washed with lysis buffer and bound material was eluted from beads with elution buffer (50 mM Tris-HCl pH 8.0, 10 mM EDTA, 1% SDS) incubating 15 min at RT and 15 min at 65°C. RIP samples along with inputs were de-crosslinked at 95°C for 10 min. Samples were then incubated with 40 μg of proteinase K (Sigma) for 4h at 37°C. RNA was recovered with a phenol-chloroform-isoamyl alcohol extraction (Thermo Fisher Scientific) followed by precipitation with sodium acetate, isopropanol and glycogen (Thermo Fisher Scientific). Precipitated RNA was digested with DNase I (Thermo Fisher Scientific) for 2h at 37°C. Purified RNA was used for reverse transcription following manufacturer’s instructions (Superscript III, Thermo Fisher Scientific) and used for qPCR as described for RT-qPCR samples. The primers used are listed in Suppl. Table 6.

### ChIP-qPCR assays

ChIP was performed as previously described (Barrales et al., 2016) with minor modifications, as follows. 100 ml of a 0.5 OD600 cell suspension was crosslinked with 1% formaldehyde (Roth) and quenched with 125 mM glycine (Sigma). Following lysis and sonication, solubilized chromatin corresponding to approx. 5-6 x 10^8^, 4 x 10^8^, 9 x 10^8^ cells was immunoprecipitated with antibodies against Pol II-S5P (25 μl supernatant, kindly provided by A. Ladurner), H3K9me2 (2 μl, Abcam 1220) and Myc-trap (10 μl, Chromotek) respectively.

RT-qPCR experiments were performed as described above, using the primers listed in Suppl. Table 6. The IP values were divided by the input and corrected for variation by normalizing to the mean of three euchromatin loci (*act1*^+^, *tef3*^+^, *ade2*^+^). The data were shown as relative to the untagged strain.

### Data and Code availability

All sequencing data that support the findings of this study have been deposited in the National Center for Biotechnology Information Gene Expression Omnibus (GEO) and are accessible through the GEO Series accession number <u>GSE174347</u>. Full code for all NGS-related workflow is available at: https://bit.ly/3wLQPqT.

## Acknowledgments

We thank members of the Braun lab as well as A. Ladurner, P. Korber, M. Halic, and M. Rougemaille for fruitful discussions during the study and critical comments on the manuscript. We thank S. Lall (Life Science Editors) for editorial assistance. We further thank A. Ladurner, M. Smolle and M. Halic for strains, plasmids and reagents; A. Yamashita; C. Brönner, M.Halic, M. Rougemaille and M. Murawska for protocols; T. Straub and T. Schauer from the BMC Bioinformatics core facility for training and counseling; G. Timinszky and J. Preisser for training and counseling in confocal microscopy; S. Stöcker for assistance in library preparation for RNA-seq analysis; and G. Schermann for genome-wide antisense transcript analysis. This study was supported by the German Research Foundation (DFG) through the collaborative research center CRC 1064 (project ID 213249687-SFB 1064) and a grant awarded to SB (BR 3511/4-1), and the Leibniz programme to IS (SI586/6-1). Additional support was provided by the Australian Government through the Australian Research Council’s Discovery Projects funding scheme to TF (project DP190100423) and by the JSPS KAKENHI Grants: JP19K06489 (to Y. Hirano); JP19K23725 (to Y. Kinugasa); JP18H05533 and JP19K22389 (to Y. Hiraoka).

## Author contributions

LMC, RRB and SB conceived the study. LMC, RRB and TvE performed RNA-seq experiments and TvE generated the RNA-seq analysis pipeline. ND, MC and LMC performed Y2H assays. MC and LMC conducted coIP experiments. LMC and SFB performed RT-qPCR experiments. All other experiments were performed by LMC. All authors analyzed the data. LMC and SB conceived and wrote the manuscript. Y. Hirano, Y. Kinugasa and Y. Hiraoka made original observations confirming independently several key findings. All authors contributed to editing.

---

**Suppl. Tables 1 and 2**: see excel files

**Suppl. Table 3.** *S. pombe* strains used in this study, related to experimental procedures

| Strain | Genotype | Source | Figure |
| --- | --- | --- | --- |
| PSB0065 | P (h+) imr1L(Ncol)::ura4+ otr1R(SphI)::ade6+ leu1-32 ura4-DS/E ade6-M210 | (Braun et al., 2011) | 1, s1, 2B, 2C, 2D, 2E, 2F, s2, s3F, 5, s5A, 6, s6D, s6E, s6F |
| PSB0906 | P (h+) leu1-32 ade6-210 ura4-DS/E imr1L(Ncol)::ura4+ otr1R(SphI)::ade6+ cen1:hphMX lem2::natMX | (Barrales et al., 2016) | 1, s1, 2B, 2C, 2D, 2E, 2F, s2, s3F, 6, s6D, s6E |
| PSB0642 | P (h-) leu1-32 ade6-210 ura4-D18 imr1L(Ncol)::ura4+ otr1R(SphI)::ade6+ mat1_m-cyhS smt0 rpl42::cyhR(sP56Q) cen1:hphMX man1::natMX | (Barrales et al., 2016) | 1F, s1D |
| PSB0640 | P (h-) leu1-32 ade6-210 ura4-D18 imr1L(Ncol)::ura4+ otr1R(SphI)::ade6+ mat1_m-cyhS smt0 rpl42::cyhR(sP56Q) cen1:hphMX ima1::natMX | (Barrales et al., 2016) | 1F, s1D |
| PSB0090 | P (h+) leu1-32 ade6-210 ura4-DS/E imr1L(Ncol)::ura4+ otr1R(SphI)::ade6+ clr4::natMX | (Braun et al., 2011) | 1G, 1H, s1E, 2D |
| PSB1480 | P (h+) leu1-32 ade6-210 ura4-DS/E imr1L(Ncol)::ura4+ otr1R(SphI)::ade6+ clr3::kanMX | (Barrales et al., 2016) | 1G, s1E |
| PSB1655 | M (h-) ura4-D18 leu1-32 ade6-M216 his7-366 swi6::natMX | This study | 1G, s1E |
| PSB1729 | P (h+) leu1-32 ade6-210 ura4-DS/E imr1L(Ncol)::ura4+ otr1R(SphI)::ade6+ clr1::kanMX | This study | 1G, s1E |
| PSB2486 (SPY418) | P h+ otr1R(SphI)::ura4+ ura4-DS/E leu1-32 ade6-M210 ago1Δ::Kan | (Halic & Moazed, 2010) | 1G, s1E, 2D |
| PSB2487 (SPY1042) | P h+ leu1-32 ade6-M210 ura4DS/E otr1(SphI)::ura4+ oriA clr2Δ::TAP-kanR | (Motamedi et al., 2008) | 1G, s1E |
| PSB1813 | P (h+) imr1L(Ncol)::ura4+ otr1R(SphI)::ade6+ leu1-32 ura4-DS/E ade6-M210 rrp6::kanMX | This study | S1C, 2B, 2C, 2D, 2E, 2F, s2A, s2I, s2J |
| PSB1849 | P (h+) imr1L(Ncol)::ura4+ otr1R(SphI)::ade6+ leu1-32 ura4-DS/E ade6-M210 rrp6::kanMX lem2::natMX | This study | 2F, s2J |
| PSB1784 | P (h+) imr1L(Ncol)::ura4+ otr1R(SphI)::ade6+ leu1-32 ura4-DS/E ade6-M210 red1::kanMX | This study | S1c, 2B, 2D, 2E, 2F, s2A, s2B, s2J |
| PSB1780 | P (h+) leu1-32 ade6-210 ura4-DS/E imr1L(Ncol)::ura4+ otr1R(SphI)::ade6+ cen1:hphMX lem2::natMX red1::kanMX | This study | 2F, s2J |
| PSB1786 | P (h+) leu1-32 ade6-M210 ura4-DS/E imr1L(Ncol)::ura4+ otr1R(SphI)::ade6+ pab2::kanMX | This study | 2D, 2F, s2J |
| PSB1781 | P (h+) leu1-32 ade6-210 ura4-DS/E imr1L(Ncol)::ura4+ otr1R(SphI)::ade6+ cen1:hphMX lem2::natMX pab2::kanMX | This study | 2F, s2J |
| PSB2061 | P (h+) imr1L(Ncol)::ura4+ otr1R(SphI)::ade6+ leu1-32 ura4-DS/E ade6-M210 air1::kanMX | This study | 2B, 2D, 2E, s2A, s2C, s6C |
| PSB1888 | P (h+) imr1L(Ncol)::ura4+ otr1R(SphI)::ade6+ leu1-32 ura4-DS/E ade6-M210 erh1::kanMX | This study | 2B, 2D, 2E, s2A, s2C, 6A |
| PSB1889 | P (h+) imr1L(Ncol)::ura4+ otr1R(SphI)::ade6+ leu1-32 ura4-DS/E ade6-M210 ccr4::kanMX | This study | 2B, 2D, 2E, s2A, s2D |
| PSB1761 | P (h+) imr1L(Ncol)::ura4+ otr1R(SphI)::ade6+ leu1-32 ura4-DS/E ade6-M210 iss10::kanMX | This study | 2D, 2E, 6B |
| PSB2651 | P (h+) imr1L(Ncol)::ura4+ otr1R(SphI)::ade6+ leu1-32 ura4-DS/E ade6-M210 Red1-9xMyc::kanMX | This study | S2H, 5D, s5A |
| PSB2653 | P (h+) leu1-32 ade6-210 ura4-DS/E imr1L(NcoI)::ura4+ otr1R(SphI)::ade6+ cen1:hphMX lem2::natMX Red1-9xMyc::kanMX | This study | S2H, 5D, s5A |
| PSB2694 | P (h+) leu1-32 ade6-210 ura4-DS/E imr1L(NcoI)::ura4+ otr1R(SphI)::ade6+ cen1:hphMX iss10::natMX Red1-9xMyc::kanMX | This study | S2H |
| PSB0374 | P (h+) leu1-32 ade6-216 ura4-D18 | Bioneer haploid deletion library | S1A, S1B, s2A |
| PSB0593 | P (h+) leu1-32 ade6-216 ura4-D18 man1::natMX | This study | S1B, s2A |
| PSB1487 | P (h+) leu1-32 ade6-216 ura4-D18 clr4::kanMX | This study | S1A, s2A |
| PSB2410 | M (h-) ura4-D18 leu1-32 ade6-M210 his7-366 Red1::6xHA::kanMX6 | This study | 3A, s6B |
| PSB2403 | P (h+) leu1-32 ade6-210 ura4 lem2::natMX Lem2::GFP::ura4+ Red1::6xHA::kanMX6 | This study | 3A |
| PSB2703 (TM15) | M (h-) lys1+::lem2p-lem2-GFP Δlem2::kanR | This study | 3C, 3D |
| PSB2704 (TM16) | M (h-) lys1+::lem2p-lem2LC-GFP Δlem2::kanR | This study | 3C, 3D |
| PSB2705 (TM17) | M (h-) lys1+::lem2p-lem2NL-GFP Δlem2::kanR | This study | 3C, 3D |
| PSB2706 (TM18) | M (h-) lys1+::lem2p-lem2L-GFP Δlem2::kanR | This study | 3C, 3D |
| PSB2707 (TM19) | M (h-) lys1+::lem2p-lem2(Δ200-307)-GFP Δlem2::kanR | This study | 3C, 3D |
| PSB2708 (TM20) | M (h-) lys1+::lem2p-GFP Δlem2::kanR | This study | 3C, 3D |
| PSB2496 | M (h-) ura4-D18 leu1-32 ade6-M210 his7-366 Red1::6xHA::kanMX6 pREP81xLem2ΔN-GFP red1-6xHA | This study | S3A |
| PSB2498 | M (h-) ura4-D18 leu1-32 ade6-M210 his7-366 Red1::6xHA::kanMX6 pREP81xLem2ΔC-GFP red1-6xHA | This study | S3A |
| PSB2494 | M (h-) ura4-D18 leu1-32 ade6-M210 his7-366 Red1::6xHA::kanMX6 pREP81x-GFP red1-6xHA | This study | S3A |
| PSB2495 | M (h-) ura4-D18 leu1-32 ade6-M210 his7-366 Red1::6xHA::kanMX6 pREP81xLem2-GFP red1-6xHA | This study | S3A |
| PSB2453 | P (h+) leu1-32 ade6-210 ura4 lem2::natMX Lem2::GFP::ura4+ | This study | S3C, s3D, s3E, 5B, s5A |
| PSB2830 | P (h+) leu1-32 ade6-216 ura4-D18 Lem2-MSC-soluble::GFP::ura4+ lem2::natMX | This study | S3E |
| PSB2831 | P (h+) leu1-32 ade6-216 ura4-D18 Lem2-MSC-soluble::GFP::ura4+ lem2::natMX | This study | S3C, s3D, s3E, s3F |
| PSB0591 | P (h+) leu1-32 ade6-216 ura4-D18 lem2::natMX | (Barrales et al., 2016) | S3F |
| PSB2840 | P (h+) ade6-M216 ura4-D18 leu1-32 Lem2::GFP::ura4+ lem2::natMX | This study | S3F |
| PSB2415 (JS71) | h90 ade6-M210 leu1 CO2::Padh1-4xU1A-Luc-natR arg1::Padh41-U1Ap-YFP-hphR | (Shichino et al. 2018) | 4B, s4A |
| PSB2416 (JS76) | h90 ade6-M210 leu1 CO2::Padh1-4xU1A-Luc-14xTTAAAC-natR arg1::Padh41-U1Ap-YFP-hphR | (Shichino et al., 2018) | 4B, s4A |
| PSB2446 | h90 ade6-M210 leu1 CO2::Padh1-4xU1A-Luc-14xTTAAAC-natR arg1::Padh41-U1Ap-YFP-hphR cut11:mCherry:HygR | This study | 4B, 4C, s4A |
| PSB2466 | h90 ade6-M210 leu1 CO2::Padh1-4xU1A-Luc-14xTTAAAC-natR arg1::Padh41-U1Ap-YFP-hphR cut11:mCherry:HygR lem2::KanMX | This study | 4B, 4C |
| PSB2846 | h90 ade6-M210 leu1 CO2::Padh1-4xU1A-Luc-14xTTAAAC-natR arg1::Padh41-U1Ap-YFP-hphR cut11:mCherry:HygR iss10::kanMX | This study | S4A |
| PSB2844 | h90 ade6-M210 leu1 CO2::Padh1-4xU1A-Luc-14xTTAAAC-natR arg1::Padh41-U1Ap-YFP-hphR cut11:mCherry:HygR red1::kanMX | This study | S4A |
| PSB1940 (YY548-13C) | h90 ade6-216 leu1-32 ura4-D18 sme2proxy[::ura4 +- kan r-lacOp] his7 +::lacI-GFP | (Ding et al., 2016) | S4C |
| PSB2524 | P (h+) leu1-32 ade6-210 ura4 Lem2::NatMX::lem2Δ | This study | 5B, s5A |
| PSB2662 | P (h+) imr1L(NcoI)::ura4+ otr1R(SphI)::ade6+ leu1-32 ura4-DS/E ade6-M210 GFP-Mmi1::natMX | This study | 5C, s5A, s6A |
| PSB2664 | P (h+) imr1L(NcoI)::ura4+ otr1R(SphI)::ade6+ leu1-32 ura4-DS/E ade6-M210 GFP-Mmi1::natMX lem2Δ::kanMX | This study | 5C, s5A, s6A |
| PSB2030 | P (h+) leu1-32 ade6-210 ura4-DS/E imr1L(NcoI)::ura4+ otr1R(SphI)::ade6+ cen1:hphMX lem2::natMX erh1::kanMX | This study | 6A |
| PSB1765 | P (h+) leu1-32 ade6-210 ura4-DS/E imr1L(NcoI)::ura4+ otr1R(SphI)::ade6+ cen1:hphMX lem2::natMX iss10::kanMX | This study | 6B |
| PSB0044 (SPY139) | h90 leu1-32 ade6-M210 ura4DS/E mat3M::ura4+ | (Bühler et al, 2007) | 6C |
| PSB0706 | h90 leu1-32 ade6-M210 ura4DS/E mat3M::ura4+ lem2::natMX | This study | 6C |
| PSB1810 | P (h+) ura4-D18 leu1-32 ade6-M210 his7-366 cut11:mCherry:hygR red1::GFP::kanMX | This study | S6A |
| PSB1837 | P (h+) ura4-D18 leu1-32 ade6-M210 his7-366 cut11:mCherry:hygR red1::GFP::kanMX lem2::natMX | This study | S6A |
| PSB2042 | P (h+) ura4-D18 leu1-32 ade6-M210 his7-366 cut11:mCherry:hygR erh1::GFP::kanMX | This study | S6A |
| PSB2094 | P (h+) ura4-D18 leu1-32 ade6-M210 his7-366 cut11:mCherry:hygR erh1::GFP::kanMX lem2::natMX | This study | S6A |
| PSB2666 | P (h+) imr1L(NcoI)::ura4+ otr1R(SphI)::ade6+ leu1-32 ura4-DS/E? ade6-M210 his7-366 GFP-Mmi1::natMX Red1::6xHA::kanMX6 | This study | S6B |
| PSB2668 | P (h+) imr1L(NcoI)::ura4+ otr1R(SphI)::ade6+ leu1-32 ura4-DS/E? ade6-M210 his7-366 GFP-Mmi1::natMX lem2Δ::kanMX Red1::6xHA::kanMX6 | This study | S6B |
| PSB2070 | P (h+) imr1L(NcoI)::ura4+ otr1R(SphI)::ade6+ leu1-32 ura4-DS/E ade6-M210 air1::kanMX lem2::natMX | This study | S6C |
| PSB2642 | P (h+) imr1L(NcoI)::ura4+ otr1R(SphI)::ade6+ leu1-32 ura4-DS/E ade6-M210 sme2::natMX | This study | S6D |
| PSB2675 | P (h+) imr1L(NcoI)::ura4+ otr1R(SphI)::ade6+ leu1-32 ura4-DS/E ade6-M210 sme2::natMX lem2::kanMX | This study | S6D |

**Suppl. Table 4.** *S. cerevisiae* strains used in this study, related to experimental procedures

| Strain | Genotype | Source | Figure |
| --- | --- | --- | --- |
| SGY137 | Y2H gold (MATa, trp1-901, leu2-3, 112, ura3-52, his3-200, gal4Δ, gal80Δ, LYS2 :: GAL1UAS–Gal1TATA–His3, GAL2UAS–Gal2TATA–Ade2 URA3 :: MEL1UAS–Mel1TATA AUR1-C MEL1) | Clontech | 3B |

**Suppl. Table 5.** Plasmids used in this study, related to experimental procedures

| Bacterial host strain | Plasmid | Insert | Marker | Source | Figure |
| --- | --- | --- | --- | --- | --- |
| ESB617 | pJK210-lem2p-Lem2-GFP-lem2t | Lem2-GFP | ampR *ura4*<sup>+</sup> | This study | 3A, s3D, s3E, s3F |
| ESB472 | pGADT7 | GAL4 AD | ampR *leu2*<sup>+</sup> | Clontech | 3B |
| ESB469 | pGBKT7 | GAL4-DBD | kanR *trp1*<sup>+</sup> | Clontech | 3B |
| ESB532 (368) | pGADT7 | GAL4 AD spRed1-1-712 | ampR *leu2*<sup>+</sup> | (Dobrev et al., in press) | 3B |
| ESB476 | pGBKT7 | GAL4-DBD Lem2-MS | kanR *trp1*<sup>+</sup> | This study | 3B |
| ESB546 (369) | pGBKT7 | GAL4-DBD spMtl1-1-1030 | kanR *trp1*<sup>+</sup> | (Dobrev et al., in press) | 3B |
| ESB379 | pREP 81xLem2ΔN-GFP | Lem2ΔN-GFP | ampR *leu2*<sup>+</sup> | (Barrales et al., 2016) | S3A |
| ESB380 | pREP 81xLem2ΔC-GFP | Lem2ΔC-GFP | ampR *leu2*<sup>+</sup> | (Barrales et al., 2016) | S3A |
| ESB382 | pREP 81xLem2-GFP | Lem2-GFP | ampR *leu2*<sup>+</sup> | (Barrales et al., 2016) | S3A |
| ESB621 | pREP 81xGFP | GFP | ampR *leu2*<sup>+</sup> | (Forsburg, 1993) | S3A |
| ESB533 (436) | pGADT7 | GAL4 AD splss10-1-551 | kanR *trp1*<sup>+</sup> | (Dobrev et al., in press) | S3B |
| ESB543 | pGADT7 | GAL4 AD spRrp6-1-777 | kanR *trp1*<sup>+</sup> | This study | S3B |
| ESB545 (438) | pGADT7 | GAL4 AD spPab2-1-166 | kanR *trp1*<sup>+</sup> | (Dobrev et al., in press) | S3B |
| ESB547 (370) | pGADT7 | GAL4 AD spMtl1-1-1030 | kanR *trp1*<sup>+</sup> | (Dobrev et al., in press) | S3B |
| ESB614 | pJK210-lem2p-NLS-MS-GFP-lem2t | NLS-MS-GFP | ampR *ura4*<sup>+</sup> | This study | S3D, s3E, s3F |
| ESB555 | pGBKT7 | GAL4-DBD Man1-MS | kanR *trp1*<sup>+</sup> | This study | S3G |

**Suppl. Table 6.**
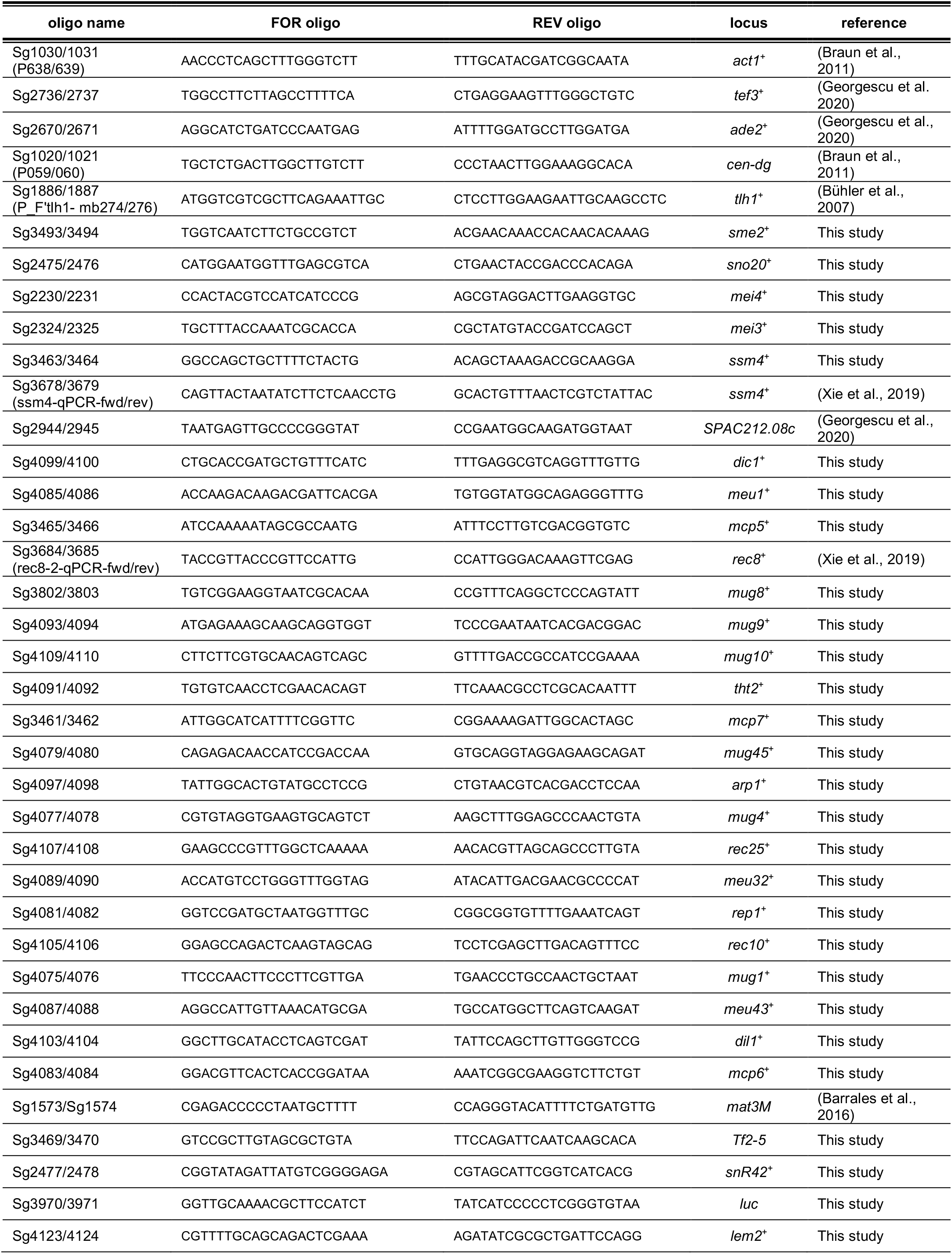
Primer sets used for RT-qPCR, ChIP-qPCR, RIP-qPCR, related to experimental procedures

**Suppl. Fig. 1:**
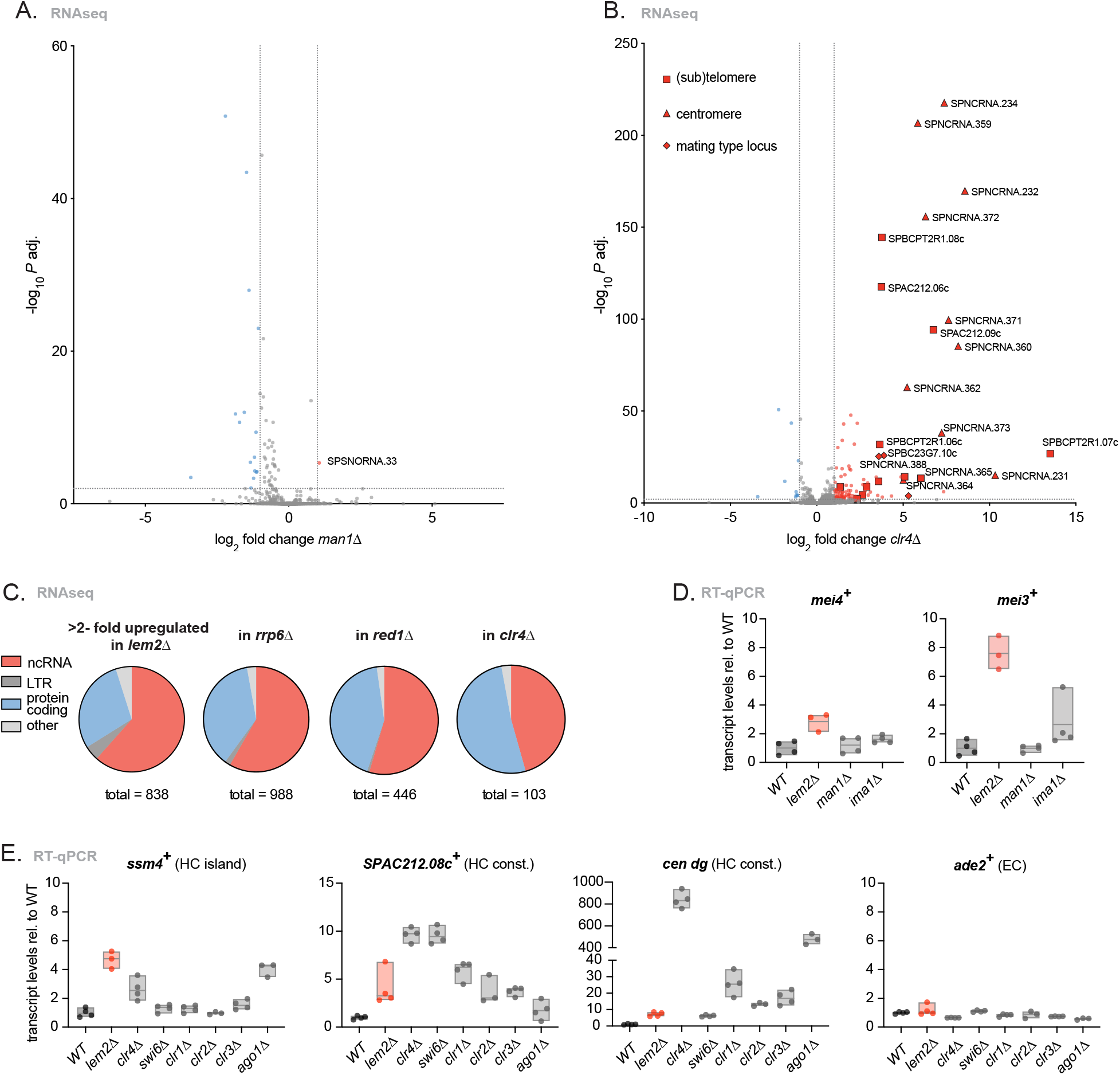
Lem2 represses non-coding RNAs and meiotic genes. **A,B.** Volcano plot depicting statistical significance (Y axis) against fold change (X axis) from the RNAseq data of *man1*Δ vs WT (A) and *clr4*Δ vs WT (B). Significant (−log_10_ *p* adj. value > 2) genes are highlighted, both upregulated (log_2_ fold change > 1 in red) and downregulated (log_2_ fold change < −1 in blue). **C.** Pie charts showing transcript feature distributions (ncRNA, LTR, protein coding, other (pseudogene, rRNA, snoRNA, snRNA, tRNA). Left: genome-wide distribution of transcript features in a WT genome. Right: transcript feature distribution of the significantly upregulated (log_2_ fold change > 1, −log_10_ *p* adj. value > 2) transcripts in the indicated mutants. **D.** Transcript levels of *mei4*^+^ and *mei3*^+^ analyzed by RT-qPCR on the indicated strains. **E.** Transcript levels of *ssm4^+^, SPAC212.08c^+^, cen dg*, and *ade2*^+^ analyzed by RT-qPCR on the indicated strains. For D and E, data was normalized to *act1*^+^ expression and it is shown relative to WT. n= 3-4 independent biological replicates. The individual replicates are shown in a floating bar plot and the line depicts the median.

**Suppl. Fig. 2:**
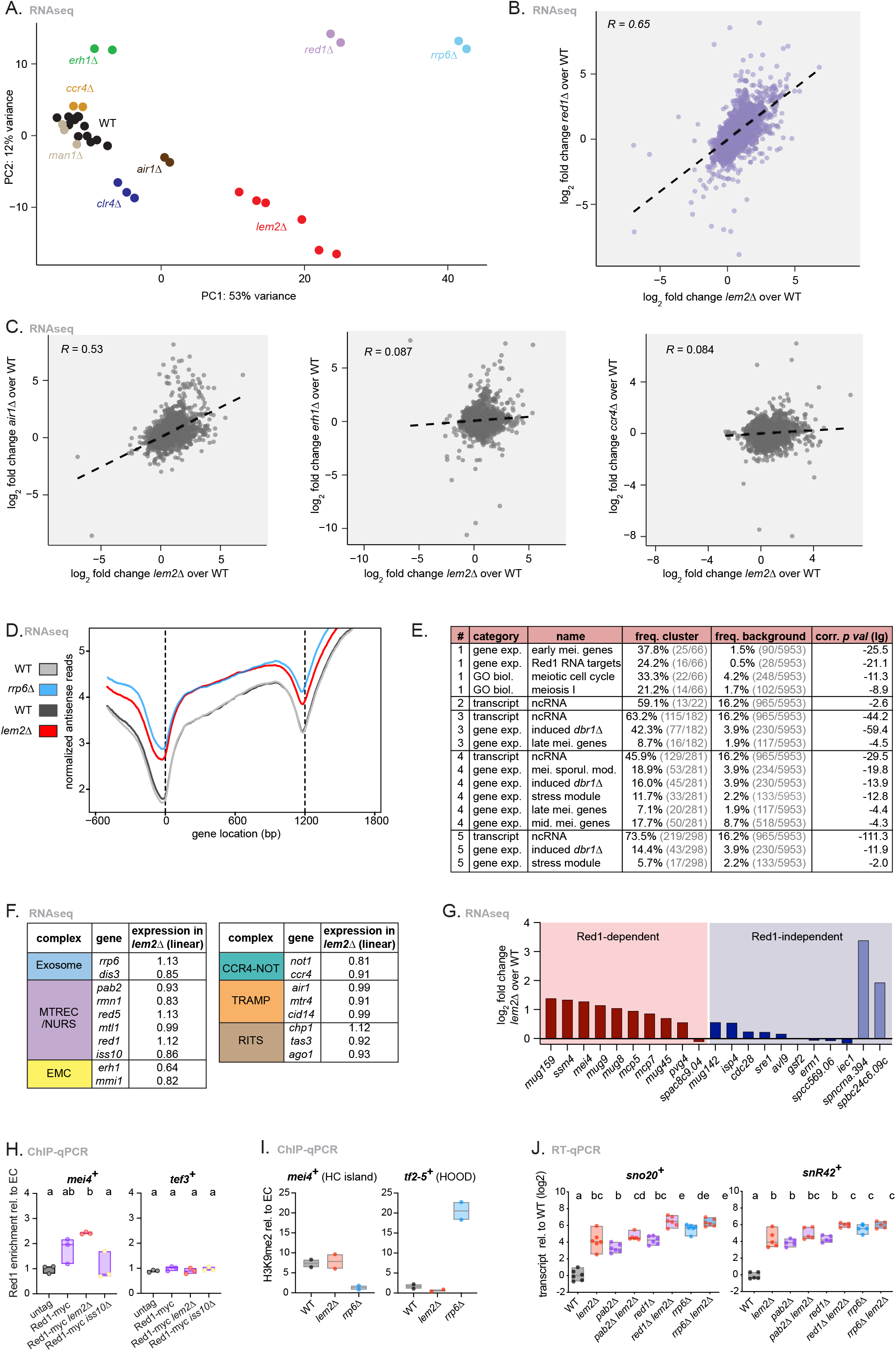
Lem2 cooperates with the nuclear exosome. **A.** PCA plot from the RNAseq libraries generated of the indicated strains. Each dot represents a biological replicate. **B, C.** Scatterplot of genome-wide log_2_ fold change expression from transcripts in *red1*Δ vs *lem2*Δ (B) or *air1*Δ vs *lem2Δ, erh1*Δ vs *lem2*Δ and *ccr4*Δ vs *lem2*Δ (C), each of them relative to WT. The linear regression line is depicted together with the Pearson correlation coefficient value (*R*). **D.** Genome-wide plot showing the log_2_ coverage of all annotated antisense mRNAs (Y axis, normalized antisense reads). All units are normalized to 1200 bp (X axis, gene location). The colors depict different strains (see legend on the left). **E.** Table with selected results from gene list enrichment analysis from the clusters (#) 1-5 from Fig. 2b. The Bähler Lab *AnGeLi* tool with FDR = 0.05 was used for this analysis (Bitton et al., 2015). Freq. = frequency; corr. = corrected; gene exp. = gene expression; GO biol. = Gene Ontology biological process; mei. = meiotic; mid. = middle. **F.** Table showing the linear expression values of multiple exosome subunits, as shown in Fig. 2a in *lem2*Δ cells. **G.** Expression levels of Red1 dependent (left, red color) and Red1 independent (right, blue color) islands in *lem2*Δ cells. Gene names are indicated below the graph. **H.** ChIP-qPCR analysis of Red1-Myc enrichment at *mei4*^+^ and *tef3*^+^ in the indicated strains. **I.** ChIP-qPCR analysis of H3K9me2 enrichment of *mei4*^+^ (island), *tf2-5*^+^ (HOOD) and *act1*^+^ in the indicated strains. **J.** *sno20*^+^ and *snR42*^+^ transcript levels quantified by RT-qPCR in the indicated strains. Data normalized to *act1*^+^ expression is shown relative to WT on a log_2_ scale. n = 6 independent biological replicates. Letters denote different groups from ANOVA and Tukey’s *post hoc* tests at *P* < 0.05. For F and G, data was retrieved from RNAseq analyses and is shown as log_2_ fold change of *lem2*Δ over WT. For H and I, the data was divided by input and normalized to euchromatin levels (*act1*^+^, *tef3*^+^, *ade2*^+^). n = 2-3 independent biological replicates. For H, I and J, the individual replicates are shown in a floating bar plot and the line depicts the median.

**Suppl. Fig. 3:**
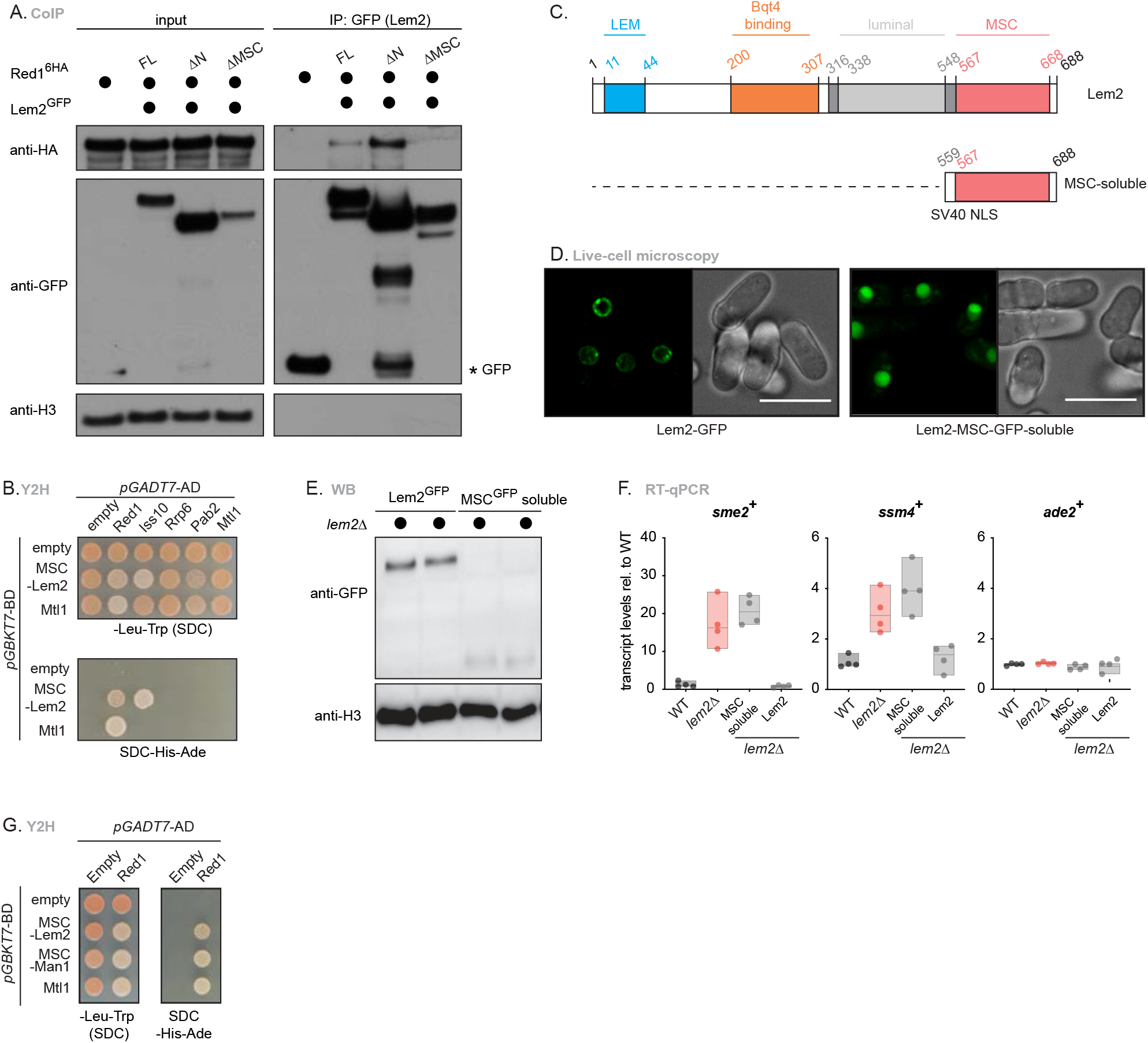
Lem2 interacts with the nuclear exosome through the MSC domain and functions in a location-dependent manner. **A.** Co-immunoprecipitation of Red1-6xHAwith Lem2-GFP WT (Lem2-GFP FL) or truncation mutants lacking the N terminal (Lem2-GFP ΔN) or MSC domain (Lem2-GFP ΔMSC). H3 served as loading control. Proteins were expressed from a *pREP81* based overexpression vector transformed in *lem2*Δ strains. **B.** Y2H analysis of Red1, Iss10, Rrp6, Pab2 and Mtl1 with MSC-Lem2 or Mtl1, grown for 3 days on medium with different auxotrophies (SDC, SDC-His-Ade). Fusions with Gal4-activating domain (*pGADT7-AD*) or Gal-4-DNA-binding domain (*pGBKT7-BD*) are shown. **C.** Schematic representation of the used Lem2 truncation constructs. Protein domains are highlighted along their amino acid length. All constructs were C-terminally GFP-tagged and inserted in the endogenous locus in *lem2*Δ background cells. **D.** Live-cell microscopy representative images of the constructs in Suppl. Fig. 3c (B). Left panels show the GFP excited channel and right panels show the corresponding bright field images. A projection of several Z-stacks with maximum intensity is shown. For some cells, Lem2-GFP is out of plane and not visible. Scale bar = 10 μm. **E.** Immunoblot of Lem2-GFP constructs in Suppl. Fig. 3c. Expression levels of two individual clones from each construct were checked. H3 served as loading control. **F.** *sme2^+^, ssm4*^+^ and *ade2*^+^ transcript levels quantified by RT-qPCR in Lem2 truncation mutants. Data was normalized to *act1*^+^ and shown relative to WT. n = 4 independent biological replicates. The individual replicates are shown in a floating bar plot and the line depicts the median. **G.** Y2H analysis of Red1 with MSC-Lem2, MSC-Man1 or Mtl1, grown for 3 days on medium with different auxotrophies (SDC, SDC-His-Ade). Fusions with Gal4-activating domain (*pGADT7*-AD) or Gal-4-DNA-binding domain (*pGBKT7*-BD) are shown.

**Suppl. Fig. 4:**
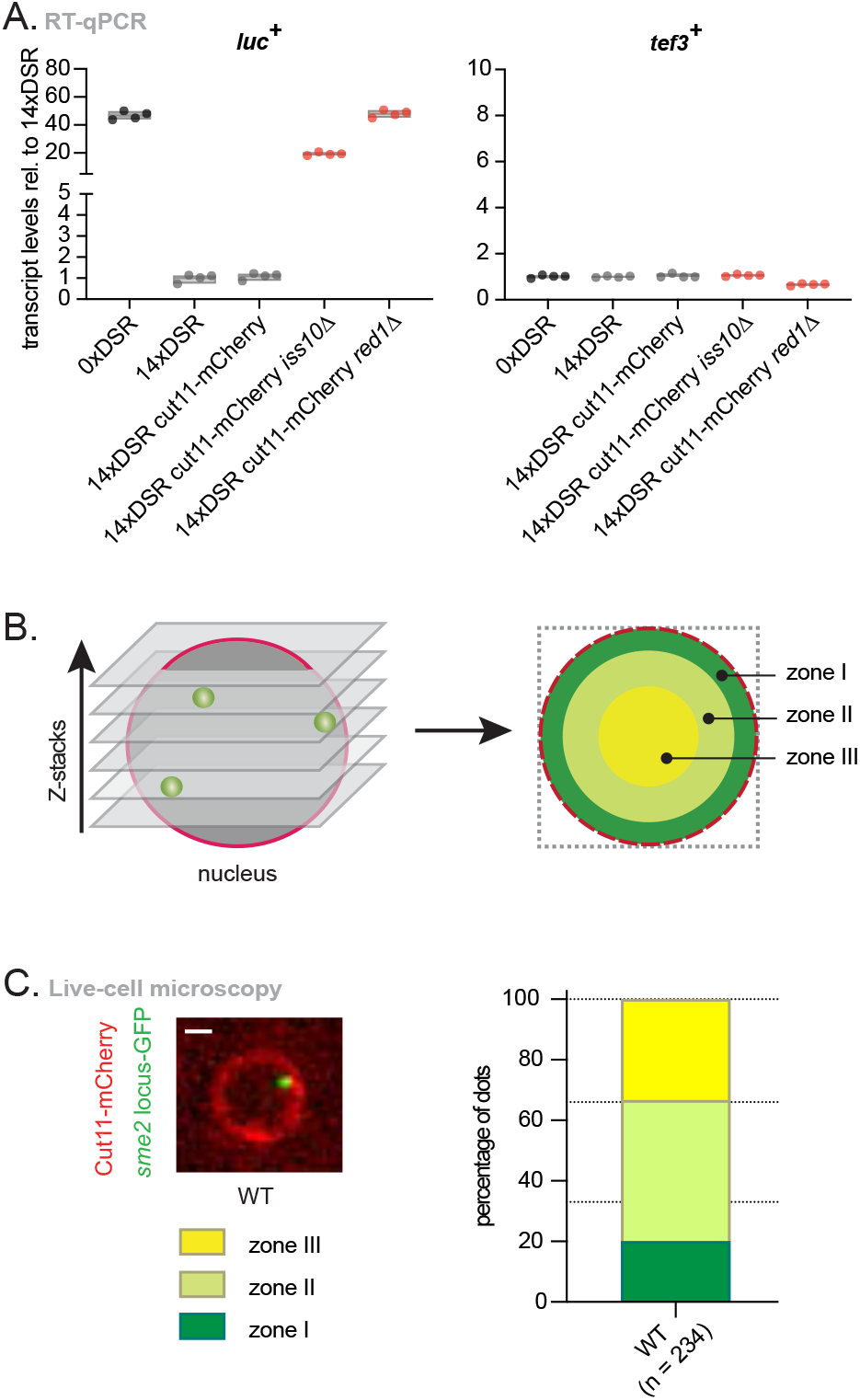
Exosome substrates localize to the nuclear periphery. **A:** Transcript levels of *luc*^+^ and *tef3*^+^ quantified by RT-qPCR on a strain encoding 0xDSR copies (0xDSR), 14xDSR copies (14xDSR), 14xDSR and the NE marker in a WT (14xDSR Cut11::mCherry), *iss10*Δ background (14xDSR Cut11::mCherry *iss10Δ)* or *red1*Δ background (14xDSR Cut11::mCherry *red1*Δ). Data was normalized to *act1*^+^ expression and shown relative to the 14xDSR strain. n = 4 independent biological replicates. The individual replicates are shown in a floating bar plot and the line depicts the median. **B:** Schematic representation of the live-cell imaging acquisition method. Z-stacks were acquired from cell nuclei detecting DSR dots and the NE (Cut11::mCherry). DSR dots localization was assigned to one out of three concentrical zones with equal surfaces within the nucleus, as measured by the distance to the NE. **C.** Top left: Live-cell microscopy representative image of the *sme2*^+^ locus (*sme2::ura4::lacOp, his7*^+^::LacI-GFP) in a WT background. Cut11-mCherry was used as a marker for the NE. A single z-stack is shown. Scale bar = 1 μm. Right: quantification of *sme2*^+^ locus location in WT and background relative to the periphery expressed in percentage of dots. n = number of cells counted in two independent experiments.

**Suppl. Fig. 5:**
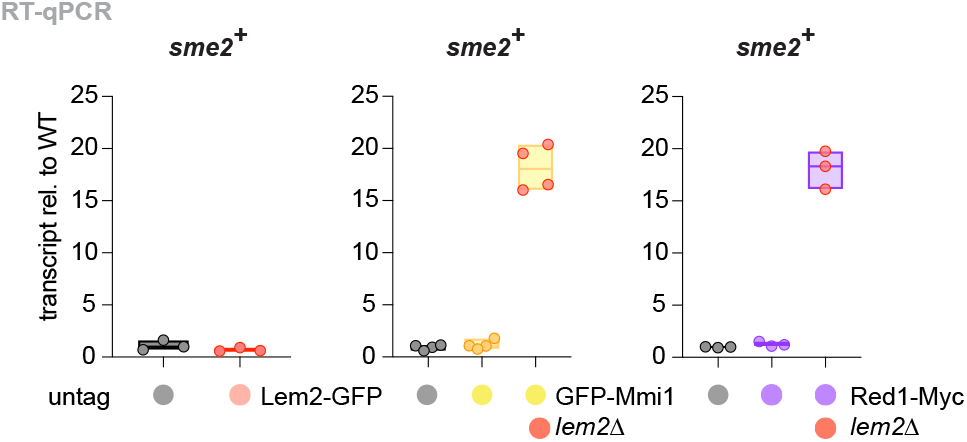
Lem2 regulates transcript binding by the exosome machinery. Transcript levels of *sme2*^+^ quantified by RT-qPCR in the input of the indicated RIP strains. Data was normalized to *act1*^+^ expression and shown relative to the untagged strain. n = 3-4 independent biological replicates. The individual replicates are shown in a floating bar plot and the line depicts the median.

**Suppl. Fig. 6:**
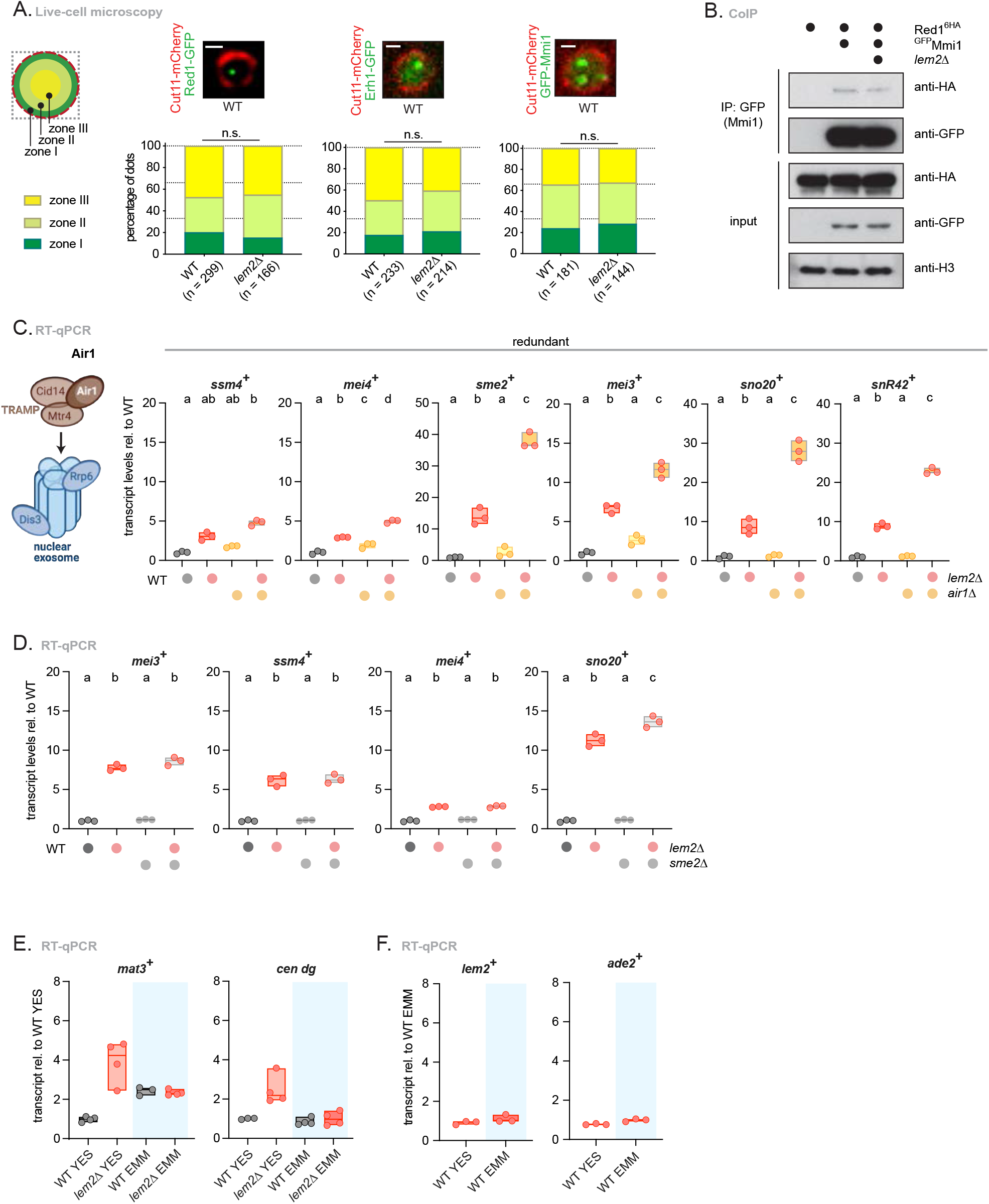
RNA regulation by Lem2 at the nuclear periphery occurs independently of exosome factors associated with nuclear foci. **A.** Top left: schematic representation of the division of a *S. pombe* nucleus in three areas with equal surfaces. These zones are called I-III depending on how close to the periphery they are. Top right: live-cell microscopy representative images of Red1-GFP, Erh1-GFP and GFP-Mmi1 in a WT or *lem2*Δ background. Cut11-mCherry was used as a marker for the NE. A single z-stack is shown. Bottom right: quantification of protein localization in WT and *lem2*Δ backgrounds relative to the periphery expressed in percentage of dots. n = number of cells counted in two independent experiments. n.s.= not significant from χ^2^ test analysis. Scale bar = 1 μm. **B.** Co-immunoprecipitation of Red1-6xHA with GFP-Mmi1 in WT or *lem2*Δ background. H3 served as loading control. Proteins were expressed from their endogenous loci. **C.** Left: scheme of the subunit that was mutated on its own or together with Lem2. Right: transcript levels of *ssm4*^+^, *mei4*^+^, *sme2*^+^, *mei3*^+^, *sno20*^+^ and *snR42*^+^ quantified by RT-qPCR in the indicated strains (*air1*Δ). Redundant or Lem2-controlled substrates are indicated above. **D.** Transcript levels of *mei3^+^, ssm4^+^, mei4*^+^ and *sno20*^+^ quantified by RT-qPCR in the indicated strains. **E.** Transcript levels of *mat3*^+^ and *cen dg* quantified by RT-qPCR in the indicated strains. **F.** Transcript levels of *lem2*^+^ and *ade2*^+^ quantified by RT-qPCR in the indicated strains. For C, D, E and F data was normalized to *act1*^+^ expression and shown relative to the WT strain (C and D), the WT in YES (D) or the WT in EMM (E). n = 3-4 independent biological replicates. Individual replicates are shown in a floating bar plot and the line depicts the median. For C and D, letters denote different groups from ANOVA and Tukey’s post hoc tests at *P* < 0.05. For E and F, blue shadowing indicates minimal media (EMM).

